# Assessing cerebellar-cortical connectivity using concurrent TMS-EEG: A Feasibility Study

**DOI:** 10.1101/2020.10.13.338350

**Authors:** Lara Fernandez, Mana Biabani, Michael Do, George M. Opie, Aron T. Hill, Michael P. Barham, Wei-Peng Teo, Linda K. Byrne, Nigel C. Rogasch, Peter G. Enticott

**Author notes:** These authors made an equal contribution to the publication (Nigel C. Rogasch, Peter G. Enticott). Corresponding Author, (L. Fernandez).

## Abstract

**Background:** Combined single-pulse transcranial magnetic stimulation (TMS) and electroencephalography (EEG) has been used to probe the features of local networks in the cerebral cortex. Here we investigate whether we can use this approach to explore long-range connections between the cerebellum and cerebral cortex.

**Objective:** To assess the feasibility of using cerebellar TMS-EEG for the exploration of cerebellar-cerebral network dynamics.

**Methods:** Ten healthy adults received single-pulse suprathreshold TMS to the cerebellum and an occipital/parietal control site with double-cone and figure-of-eight coils while cerebral activity was recorded. A multisensory electrical control condition was used to simulate the sensation of the double-cone coil at the cerebellar site. Two cleaning pipelines were compared, and the spatiotemporal relationships of the EEG output between conditions were examined at sensor and source levels.

**Results:** Cerebellar stimulation with the double-cone coil resulted in large artefacts in the EEG trace. The addition of SOUND filtering to the cleaning pipeline improved the signal such that further analyses could be undertaken. The cortical potentials evoked by the active TMS conditions showed strong relationships with the responses to the multisensory control condition after ~50 ms. A distinct parietal component at ~42 ms was found following cerebellar double-cone stimulation.

**Conclusions:** Cerebellar double-cone stimulation produces large artefacts in the EEG. Cerebellar-specific responses could not be reliably differentiated from sensory evoked potentials after ~50 ms. While evoked potentials differed across conditions at early latencies, it is unclear as to whether these represented TMS-related network activation of the cerebellarthalamocortical tract, or whether components were dominated by sensory contamination and/or coil-driven artefacts. Further work will be required to clarify the specific contribution of cerebellar-cortical connectivity to the observed early latency signals.

## Introduction

The emerging understanding of the role of the cerebellum in higher cognitive functions such as theory of mind [1] and social sequencing [2, 3] has sparked a surge of interest toward cerebellar-cortical connectivity. Structural and functional associations between cerebellar and cerebral sites have been revealed using techniques with high spatial resolution, such as viral tracing in non-human primates [4], and magnetic resonance imaging (MRI) [5, 6]. One means of exploring the temporal features of a neural network is via transcranial magnetic stimulation (TMS). Single-pulse TMS produces a time-varying electric field that can directly excite neural populations at the site of stimulation. A bifocal TMS paradigm involving primary motor cortex (M1) stimulation preceded by cerebellar stimulation with a double-cone (DC) coil (which allows deeper pulse penetration than flat coils), has been used to probe the conduction times of cerebellar-to-M1 connectivity [7–9]. Using this approach, motor evoked potentials (MEPs) recorded from the periphery are inhibited at short (5-7 ms) interstimulus intervals, revealing a rapid inhibitory influence of cerebellar stimulation on motor cortex excitability that is thought to be mediated by cerebellarthalamocortical connections [7, 9].

However, the temporal features of TMS-induced network-based transfer from locally activated Purkinje cells in the cerebellar cortex to cerebral regions via the cerebellarthalamocortical tract are not well characterised. Electroencephalography (EEG) has increasingly been used to quantify the direct cortical response to focal TMS, particularly in regions not amenable to peripheral motor assessment [10–16]. While a small number of studies have reported changes in cortical oscillations and TMS-evoked potentials (TEPs) following single-pulse cerebellar TMS, they did not always include appropriate controls to account for the large muscle and sensory artefacts that accompany the TMS pulse (see [17] for review). Indeed, muscle and sensory artefacts are well documented in the TMS-EEG literature, and deemed notoriously difficult to separate from TEPs due to substantial spatial and temporal overlaps [12–14, 18–21]. The non-linearity of these relationships necessitates the use of sophisticated methods to extract the salient features of the TMS-evoked signal. This is problematic for cerebellar TMS-EEG, as the DC coil emits a broad electromagnetic field that, when placed near the base of the skull, produces visible activation of neck, jaw, and scalp muscles in the healthy adult subject. Furthermore, studies using MEPs have shown that to sufficiently target Purkinje cells in anterior portions of the cerebellar cortex, stimuli should be delivered by a DC coil at upwards of 60% of the maximum stimulator output (MSO), or higher when using planar coils [8, 22, 23]. The resultant large-amplitude muscle and sensory activity will inevitably be recorded by the EEG and confound the underlying signal of interest.

The current study assessed the feasibility of obtaining TEPs from single-pulse cerebellar TMS using a DC coil combined with concurrent EEG. An intensity of 60 %MSO over the right cerebellar hemisphere was used to ensure activation of the cerebellar cortex [22]. To control for somatosensory and auditory components introduced by the TMS pulse in the absence of cerebellar activation, we implemented a realistic multisensory control condition including electrical stimulation of the target region, and associated coil click. This was designed to replicate the muscle, somatosensory and auditory response to the DC coil when placed over the right cerebellar hemisphere. Additional control conditions were included to assess coil-type and site-specific effects, comprising cerebellar stimulation with a figure-of-eight (F8) coil, and occipital/parietal stimulation with a DC and F8 coil. We assessed similarities and differences between conditions using raw and processed signals, and explored the effects of using different cleaning pipelines on the raw data.

## Method

### Participants

Ten healthy participants were recruited via online and physical advertisements (6 female, mean age = 21.40 years, SD = 2.76). All provided fully informed written consent to participate in a single session and received a $20 AUD department store voucher as reimbursement. Participants were right-handed according to the Edinburgh Handedness Inventory [24] and were screened for any TMS contraindications using standard exclusionary criteria [25]. Participants were also screened for coordination deficits according to the Adult DCD/Dyspraxia Checklist (ADC), as a high score may indicate cerebellar dysfunction. The study was approved by the Deakin University Human Research Ethics Committee.

### Experimental Design

In a single session, participants underwent five different single-pulse stimulation conditions and a resting-state condition. The order of the conditions was counterbalanced across participants. The four active TMS conditions involved application of stimulation to either the right cerebellar hemisphere, or an occipital/parietal control location to test site specificity, using both DC and F8 coils. To activate the cerebellum, a stimulation site that was 3 cm right of the midline and level with the inion was used [7, 9] (Figure 1,Supplementary Figure S1). For occipital/parietal control stimulation, a site was chosen that was sufficiently posterior to produce a similar sensation to the cerebellar location, yet anterior enough to exclude cerebellar tissue. To avoid eliciting confounding phenomena such as phosphenes and other visual disturbances when stimulating the occipital lobe, a standard procedure was applied for both coils prior to EEG recording [26] (Supplementary Material). In the current study, all active control stimulation was applied over electrode PO4, as no participant reported phosphenes at this location.

**Figure 1:**
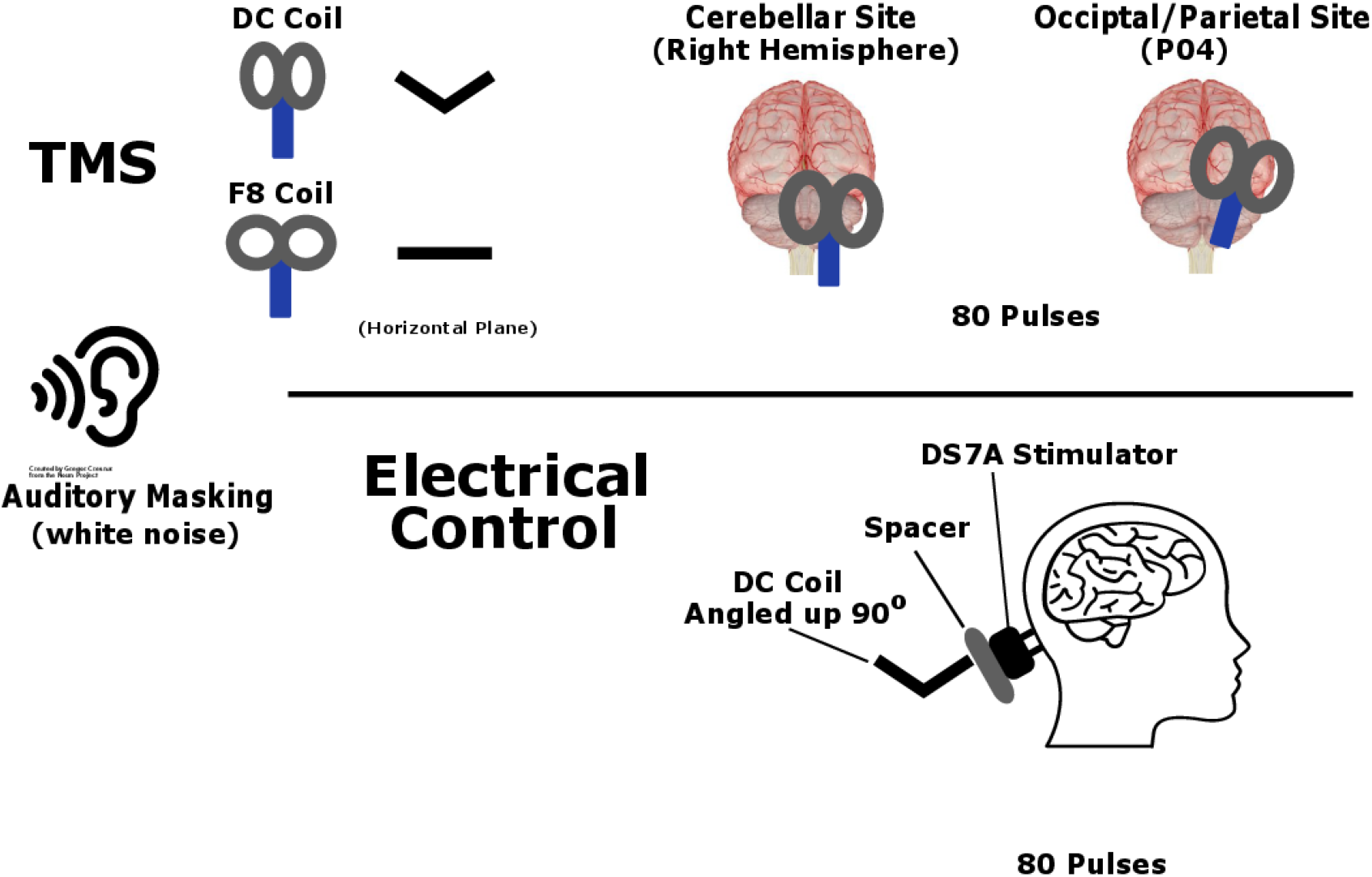
Experimental set-up. Eighty TMS stimuli were delivered to the right cerebellar hemisphere via a DC coil. Stimuli were also delivered to the cerebellum with an F8 coil, and to an occipital/parietal site with both coil types to act as active controls. A multisensory control delivered electrical stimuli to the cerebellar site via a DS7A stimulator, while simultaneously discharging the DC coil next to the head. Auditory masking was used in all conditions.

A thin layer of foam (5-10 mm) was placed over the face of each coil to minimise contact with the EEG electrodes and dampen vibrations during discharge [27]. A realistic control condition was implemented using cutaneous electrical stimulation with concurrent TMS coil discharge next to the head to simulate the ‘click’ sound of the coil [11, 12]. To minimise contamination of the EEG trace by auditory-evoked artefacts, white noise was played through noise-cancelling earphones (Compumedics Earphone Insert 10 Ω ¼ stereo, used with Etymotic Research foam ear inserts), and individually adjusted to a threshold chosen by participants to minimise the sound of the coil ‘click’ while also minimising discomfort from high volume playback.

### Sensory Rating Scales

To measure the extent to which TMS-related somatosensory and auditory stimuli may have influenced EEG outcomes for each coil/location pair, participants completed four visual analogue scales (VAS) immediately following each condition. To quantify subjective levels of discomfort and degree of muscular ‘twitch’ induced by the TMS pulses, participants were asked to indicate on a continuous 11-point scale (0 – minimum, 10 - maximum), ‘How uncomfortable were the TMS pulses?’, ‘If painful, how intense was the pain from the pulse?’, and ‘If there were any twitches, how strong were the muscle twitches from the TMS pulse?’. A fourth scale assessed the perceived loudness of the coil ‘click’ via, ‘How well did the white noise cover the sound of the TMS pulse?’.

### EEG Recording

EEG was acquired with a TMS-compatible SynAmps RT system (Compumedics Ltd, Melbourne representative, Australia). Recordings were obtained from 60 low-profile Ag/AgCl sintered electrodes with c-slits to reduced eddy current build-up, and embedded in a plastic cap (EASYCAP, Germany). Electrodes were positioned according to the 10-20 international system, with FCz as reference, and AFz as ground. Four free electrodes were used to monitor eye movement. Eye channels and TP9 and TP10 electrodes were removed offline before analysis as these frequently did not sit flush against the participant’s scalp. Signals were low-pass filtered (4 kHz) and digitised (10 kHz) online via Curry 7 software (Neuroscan, Compumedics, Australia). Skin-electrode impedance levels were monitored throughout the session and kept below 5 kΩ.

### Transcranial Magnetic Stimulation

For the TMS-EEG trials, TMS was administered via a 110 mm Magstim DC (Ref. 4610-00) and 70 mm F8 coil (Ref. 3190-00). Coils were connected to a Magstim 200^2^ unit (Magstim, UK) to deliver monophasic pulses. 80 stimuli were delivered in five second intervals (±10%) using a PowerLab 4/35 system (ADInstruments, Dunedin, NZ) and LabChart8 software (ADInstruments, Dunedin, NZ) to either cerebellar or occipital/parietal sites. Stimuli were delivered by the DC coil at 60 %MSO, as this intensity has been previously found to be a lower bound for activating the anterior cerebellum using the same stimulator and coil [22]. To account for the reduced depth-range of the F8 coil [28], stimuli were delivered at 90 %MSO by the F8 coil to ensure that the electromagnetic field reached cerebellar tissue [29] (and personal communication with authors). These intensities were also used at the occipital-parietal site for consistency. Coils were held with handle pointing downwards in order to elicit an upward current in the brain [7, 9].

### Multi-sensory Control Condition

A realistic sensory control condition was implemented via a DS7A electrical stimulator producing a 200 μs square pulse, with maximum compliance voltage of 200 V (Digitimer Ltd., Ft Lauderdale, Florida, USA) [11, 12]. Bipolar electrodes (25 mm apart) were soaked in saline solution attached to a 36 mm plastic frame. On the participant’s right side, hair was parted in order to maintain a good contact with the skin, and electrodes were pressed against the scalp to elicit a twitch sensation in neck muscles resembling that of the DC coil at the cerebellar site. This typically occurred when electrodes were placed at the base of the skull over proximal neck muscles, avoiding the spine. The stimulating electrodes were always situated below the level of the cap, and not over the cerebellum, as this can result in activation of neural tissue at sufficient intensities [30]. The site and level of current delivered were individualised according to participant feedback and visual observation of muscle activity. To mimic the auditory sensation of TMS from both air and bone conduction, the DC coil was positioned next to the electrical stimulation site and discharged in synchrony with the electrical stimulus at a frequency identical to the active TMS conditions using Powerlab and LabChart software. To ensure that the field from the TMS coil resulted in no appreciable neurological effect, the DC coil was angled upwards by 90° (lead pointing down, perpendicular to the ground) and separated from the scalp by a solid block of hardened modelling clay (30 mm diameter). The intensity of the TMS pulse was increased by 5 %MSO to account for the increased distance between coil and scalp relative to the active TMS condition [12].

### EEG Pre-Processing

EEG data were analysed using custom scripts implemented in MATLAB (R2015b, The Mathworks, USA), with EEGLAB [31] and TESA [16] toolboxes. Two cleaning and analysis pipelines were used on the raw data. The first pipeline was based on a previously described method tailored for M1 stimulation with a flat F8 coil [16] (Supplementary Material). A second pipeline included source-utilized noise-discarding (SOUND) filtering [32, 33] prior to the first ICA step to characterise and suppress large artefacts in individual channels and trials (Supplementary Material). A SOUND regularisation parameter of 0.1 was used, as this was suggested in the original publication to provide the best compromise between improving the SNR and overcorrection of the signal [33]. Raw and cleaned data are available [34]. The proportion (of total) of trials and channels rejected for each condition following each pipeline is shown in Table S1.

### Suppression of Sensory Evoked Potentials

To further assess whether any TEP peaks were independent from peaks following sensory stimulation, we applied signal-space-projection-source-informed-reconstruction (SSP-SIR) [21]. SSP-SIR is a spatial filtering method designed to estimate the artefact-free neural signal acquired with EEG using known artefact properties (e.g. frequency or spatiotemporal profile) which differ to the signal of interest. However, where the frequency properties of standard muscle artefacts were used in the original implementation [21], we took the approach outlined in [14], and used the signal obtained from the multisensory control condition to determine the artefactual subspace. As described in this publication, we removed the *k* components that explained more than 90% of variance in the electrical stimulation trials from the cerebellar-DC data.

### Source Analysis

Source estimates for the average-referenced cleaned and SSP-SIR signals were derived using the Brainstorm software [35]. A common head model was computed using a three-layer symmetric Boundary Element Method (BEM, OpenMEEG) using the default anatomy (ICBM152) and 1922 vertices per layer. Conductivities were set to the default values (scalp = 1, skull = 0.0125, brain = 1). Each individual’s datasets were then imported into Brainstorm and averaged over epochs to obtain the time-locked trace for each condition. Noise covariance matrices were derived from the pre-stimulus recordings (−1 s to −0.001 s). Cortical sources were estimated using minimum norm estimation and current source density (depth-weighting order = 0.5, regularisation parameter signal-to-noise ratio = 3.0), with dipoles assumed to be tangential to the surface. Estimates were converted to z-scores, and a weighted average was then performed over participants to form the grand average of a condition. Source estimates were smoothed with a kernel of size 3 mm for display.

### Statistical Analysis

All statistical analyses of TMS-EEG data were performed in MATLAB (R2015b) or SPSS (version 26, IBM Corporation, Armonk, NY). Scores from the VAS were analysed in SPSS. As the data for each sensory scale were normally distributed (KS-test with Liliefors correction was non-significant at p<0.01), a two-by-two (site, TMS coil-type) repeated-measures factorial ANOVA (Bonferroni corrected for p<0.05) was conducted for each scale across TMS stimulation conditions. Repeated-measures t-tests were conducted to assess differences between TMS condition and the multisensory control. To assess channel recovery times for different coil-site conditions, the time (ms) for each channel to return to a magnitude at or below ± 150 μV was found [19] and statistical comparisons performed using non-parametric tests (Wilcoxon Signed Rank), as data were not normally distributed (Supplementary Material). ICA performance (variance and components removed) was also assessed non-parametrically (Wilcoxon Signed Rank).

Spearman’s rank correlation analyses were used to explore statistical relationships between the cleaned signals from different conditions in both spatial (i.e., across electrodes for each time point) and temporal domains (i.e., across time points for each electrode). For the temporal analyses, latencies were grouped into early (20-60 ms), mid (60-180 ms) and late (180-280 ms) time ranges [14]. To analyse these correlation values at the group level, a procedure based on Fisher’s transform for dependent samples was used to convert the correlation statistics to z-values [14, 36, 37]. Statistical significance was determined using a one-sample permutation test assessing the hypothesis that each z-score was greater than zero (i.e., positive correlation). To control the family-wise error rate for multiple comparisons (time-points for the spatial correlations, and channels for the temporal correlations), we used the tmax method described in [38]. The z-values were subsequently transformed back into their original form (ρ) for display. To further compare the spatio-temporal voltage profiles of the electrical control against the active TMS conditions and across sites and coil-types, cluster-based permutation tests (paired-sample t-tests) were applied via the Fieldtrip toolbox [39] to the cleaned and average-referenced data. Here, clusters were defined as at least two adjacent electrodes exceeding p-value < 0.05 at each time-point. The critical alpha level for Monte-Carlo p-values was set at p<0.025 (i.e., two-tailed test), using 5000 iterations.

An additional spatial filter, the surface Laplacian, was also applied to the cleaned data to aid comparisons by improving topographical localisation by minimising volume conduction [40]. Computations were based on the spherical spline method described in Perrin, Pernier (41). The Legendre polynomial order was set to 50, and smoothing parameter (lambda) set to 10^−5^. Permutation tests and correlation analyses were subsequently applied to the spatially filtered data, and presented separately to the average-referenced data.

## Results

All sessions were run to completion, and no serious adverse side effects were observed or reported by participants. However, participants frequently stated that the stimuli from TMS and electrical stimulation were uncomfortable. One participant reported poor tolerance to the electrical stimuli (multisensory control), necessitating the use of a low current (9 mA). As this did not produce a sufficient muscle response, these trials were excluded from analyses (i.e. participant 6 removed from all comparisons using the multisensory control). In the remaining nine participants, the current ranged from 20.5 mA to 40 mA (mean 32.28 mA, SD 7.45). In all cases, a distinct muscle twitch (jaw, scalp and neck) was reported by the participant and visually observed by the researchers both during pulse administration and in the resultant EEG trace. The currents used were higher than those reported in Conde, Tomasevic (12) for the M1. However, as the stimulating electrodes were deliberately placed over the muscles and not directly over the cerebellum, the possibility of cerebellar stimulation by the electrical stimulus itself was minimised.

### Sensory rating scales

The descriptive statistics for the VAS are shown in Table S2. Repeated measures factorial ANOVAs were performed for each scale separately to assess differences between the active TMS conditions using stimulation site (cerebellum, occipital/parietal) and TMS coil type as factors (DC, F8). We could not find evidence for a difference between the pain and auditory masking scales between stimulation conditions (all p-values > 0.05, Bonferroni correction). A significant main effect of coil-type was found for the discomfort scale (*F*(1,9)=16.89, p<.01, ηp^2^=.65), where the DC coil was rated as more uncomfortable than the F8 at both stimulation sites. A significant main effect of coil-type was also found for the twitch scale, (*F*(1,9)=17.60, p<.01, ηp^2^=.66), and a main effect of site (*F*(1,9)=8.49, p=0.02, ηp^2^=.49), with no interaction effect. The twitch sensation at the cerebellar site was greater than at the occipital/parietal site for both coil types, where the DC coil produced an overall greater twitch response than the F8. When contrasting the DC conditions with the multisensory control, repeated-measures t-tests between the cerebellar (CB)-DC and control found a significant effect for the twitch scale only (*t*(9)=2.50, p=.03, d=.62), where the sensation following DC stimulation (mean=5.12, SE=.72) was greater than following electrical stimulation (mean=3.62, SE=.77), 95% CI [.14, 2.86]. We found no evidence for a difference between the occipital/parietal (OP)-DC condition and multisensory control (all p-values > 0.1).

Although underpowered due to low sample-size, these results suggest that the DC coil was more uncomfortable than the F8, particularly at the cerebellar site, which was also accompanied by a stronger twitch sensation (Figure 2). Indeed, participants were sometimes observed moving away from the coil during DC pulse administration (coil position was adjusted online by an experimenter to ensure good contact with the scalp). The CB-DC condition also produced a greater twitch response than the multisensory control. Notably, electrical stimulation was numerically more painful than TMS, although less so when comparing against the CB-DC condition. The electrical stimulus was also frequently referred to as ‘unpleasant’, which was largely attributed to the direct ‘sharp’ sensation of the pulse on the skin, and resultant focal muscle twitch. As intensities were constrained by the maximum current tolerated by the subject, this posed a limitation to the effectiveness of the multisensory control condition.

**Figure 2:**
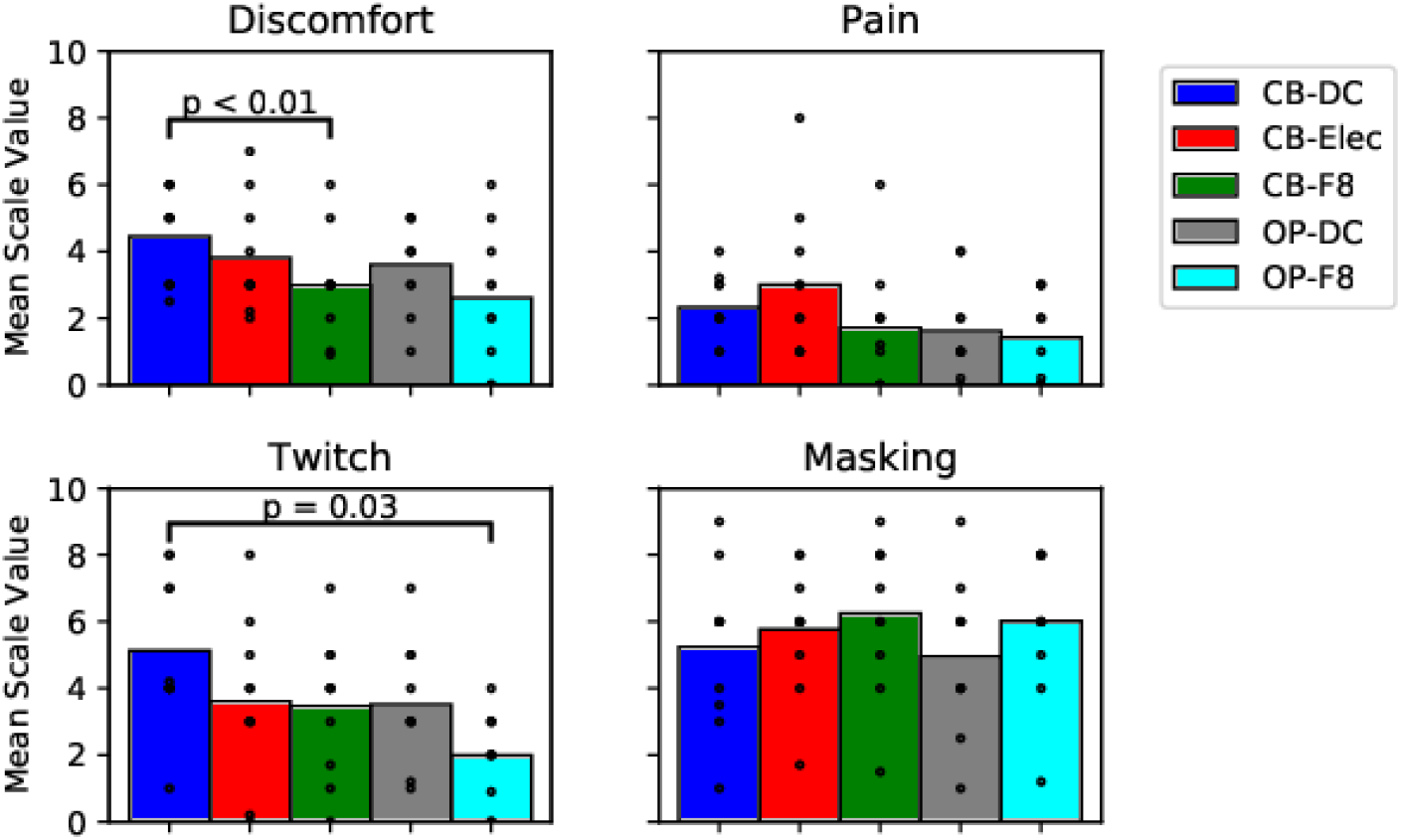
Visual Analog Scales. Mean scale values for each condition (site-coil type) averaged over participants, with raw data superimposed. CB - cerebellar site, OP - occipital/parietal site, DC - double-cone coil, F8 – figure of 8 coil, Elec – electrical (multisensory control) stimulation. Continuous scales were between 0-10, and measured the discomfort of stimulation (discomfort scale), associated pain (pain scale), twitch sensation accompanying the pulse (twitch scale), and degree to which white noise masked the sound of the TMS coil (masking scale).

All conditions were generally well matched in auditory masking. However, auditory masking was not overly successful in any condition, with nearly all participants indicating only moderate masking efficacy.

### Raw Data and Cleaning Pipelines

Two datasets were removed due to poor signal quality (CB-F8 for participants 5 and 6, thus only eight participants were used in comparative analyses for this condition). The remaining raw traces showed similar artefact profiles across all stimulation conditions. All stimulation conditions resulted in a pulse artefact that peaked within the first 2 ms (Figure 3A). Additional peaks consistent with muscle activity were also observed ~5-8 ms post-pulse (~1-2 mV in amplitude, Figure 3B). Both TMS pulse and muscle-related artefacts were removed from the data and interpolated (−2 to 10 ms).

**Figure 3:**
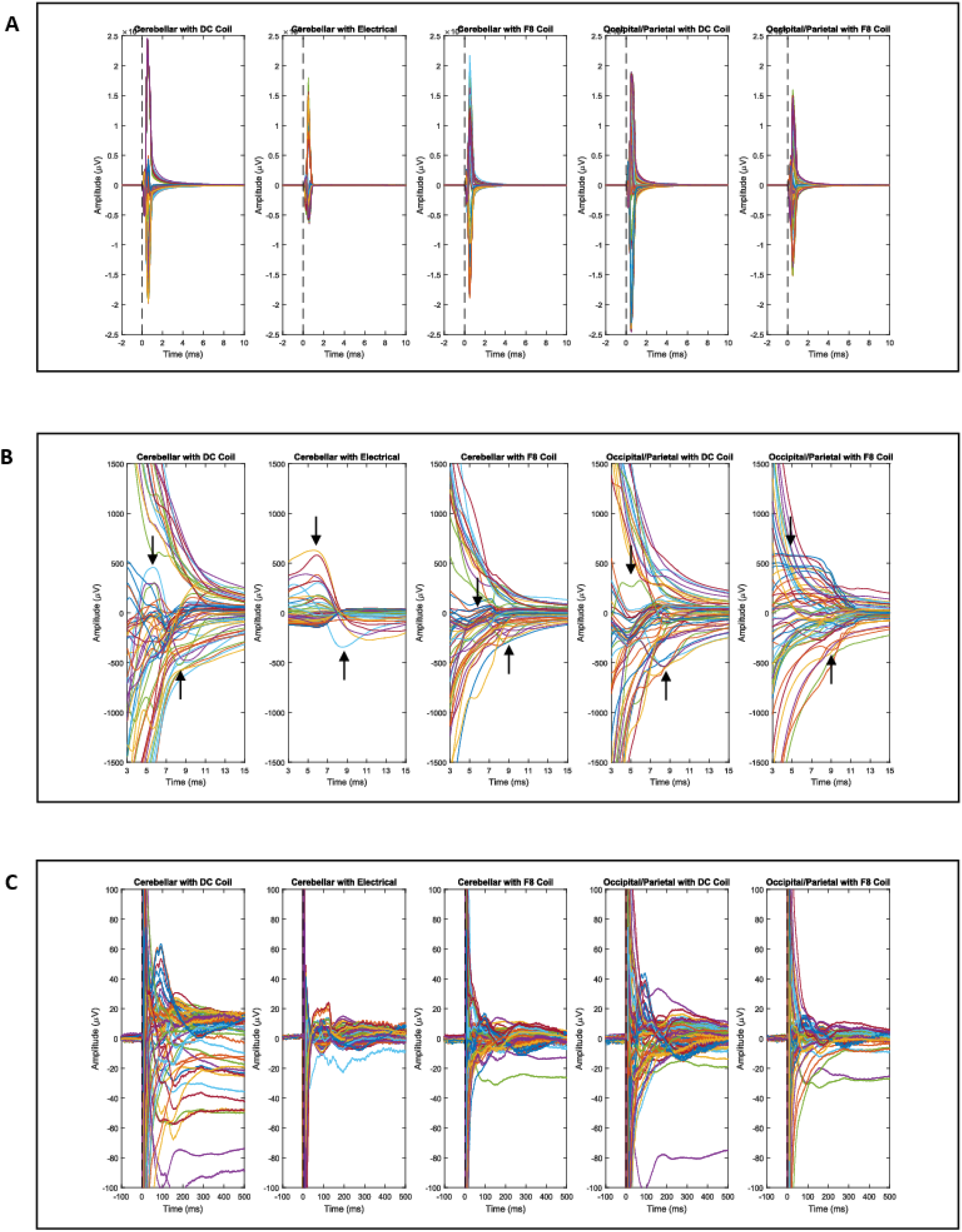
Raw grand averaged timeseries for each condition and each of the 60 electrodes re-referenced to the common average. A. Signals zoomed out to maximum global range to see pulse-related artefacts. Broken vertical line shows pulse onset. B. Peaks related to muscle artefact at 5 ms and 8 ms post-pulse (arrows). C. Signals zoomed in to within ± 100 μV showing decay artefacts. Pulse onset at 0 ms. Larger deviations from baseline are noticeable with the DC coil relative to the F8 coil, regardless of stimulation site, although stimulation of the cerebellum with the DC coil produced the highest deviations.

Large voltage offsets in the EEG trace were present following all stimulation conditions, with recovery taking up to several hundred milliseconds. Offset amplitudes were generally greatest in channels closest to the site of stimulation (Figure 3C,S2), and particularly large and widespread following cerebellar stimulation with the DC coil (Table S3-5). However, voltage ranges did not differ significantly between the DC and F8 coil at the cerebellar site (p = .40), suggesting that the signal contained a large degree of artefact when TMS was applied with either coil to the cerebellum. Furthermore, the voltage range from the CB-DC condition was significantly higher than that for the multisensory control (p = .01), indicating that the sensory control condition may not have optimally triggered all TMS-evoked sensory components present in the CB-DC condition.

Signal recovery times were greater for the DC coil at the cerebellar site relative to the F8 coil (p = .03), and the DC coil also produced longer recovery times at the cerebellar versus occipital/parietal site (p = .01) (Table S4, S5). The presence of such large decay artefacts can prove problematic for blind-source separation algorithms such as ICA, which are sensitive to low frequency drifts in the data [42] and activity that is highly time-locked to the neural signal [43]. Indeed, decay artefacts dominated many independent components following the first and second round of FastICA (Figure 4). As a result, the common two-stage ICA cleaning pipeline was suboptimal for these data sets. This pipeline introduced large baseline offsets likely due to interactions with temporal filters following insufficient removal of high amplitude artefacts, and failed to suppress decay artefacts in some electrodes.

**Figure 4:**
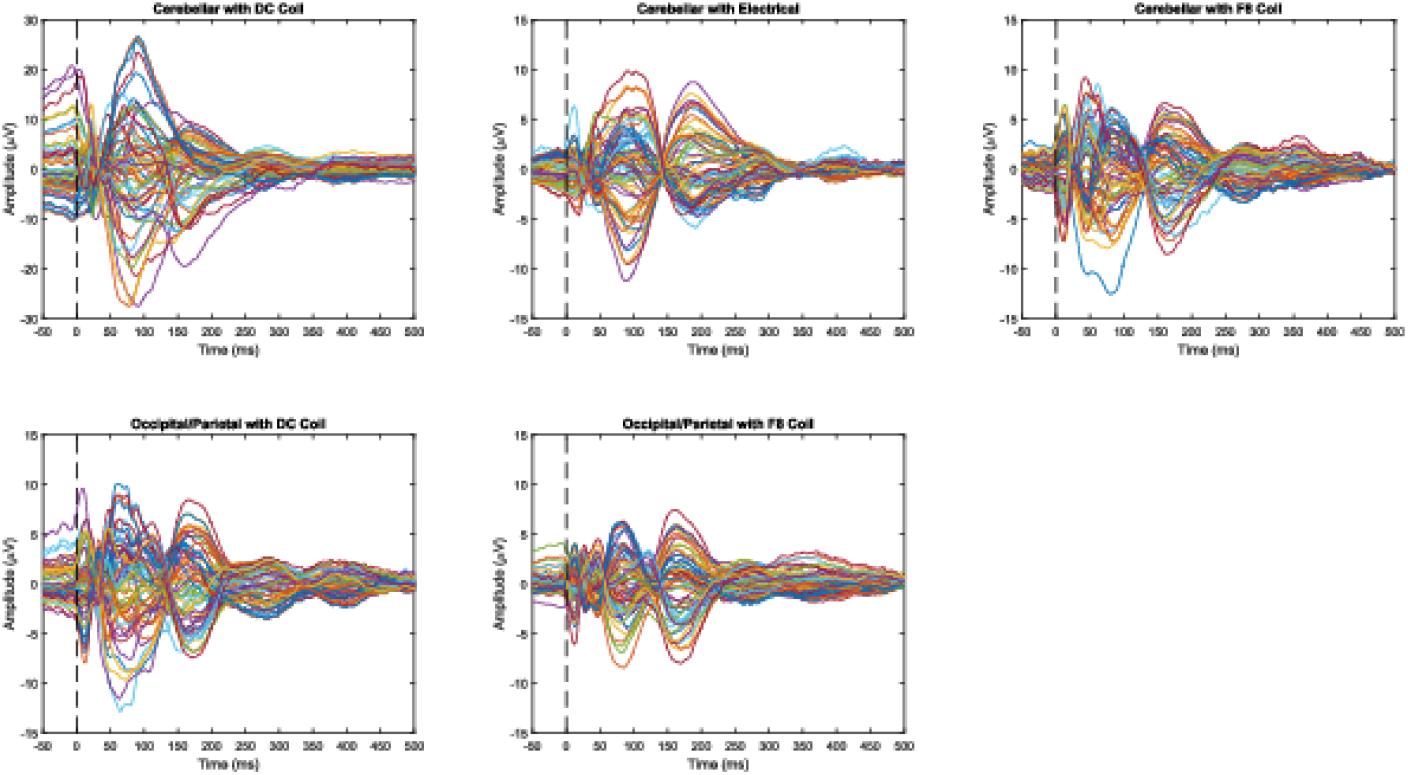
Grand-average signals following cleaning using the Original pipeline, for each condition. Broken vertical line shows pulse onset.

Given the high levels of artefacts resulting from DC stimulation, we assessed whether including SOUND prior to ICA improved the cleaning pipeline performance by suppressing decay artefacts in highly affected channels. SOUND filtering reduced signal noise in the pre-stimulation and early (< 60 ms) periods (i.e., resulted in less ‘ringing’ artefact from bandpass filtering), and generally produced clearer peaks relative to the initial pipeline (Figure 5A). This was particularly apparent in the active cerebellar conditions and was also reflected in the spatio-temporal voltage distributions for each condition (Figure 5B).

**Figure 5:**
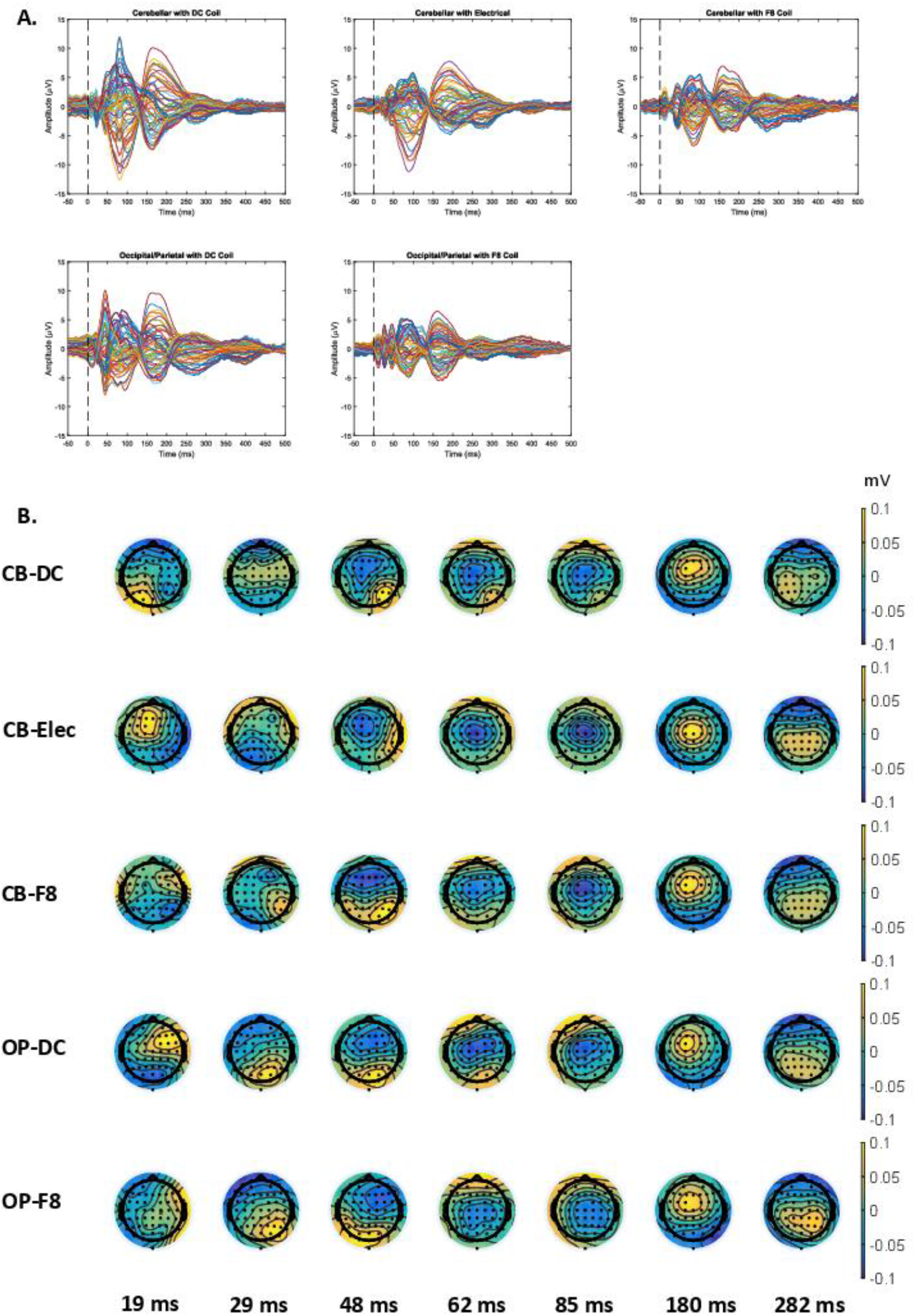
A. Grand-average of signals following data cleaning using the SOUND pipeline, for each condition. Broken vertical line shows pulse onset. B. Spatiotemporal distribution of grand average signal for each condition following cleaning using the SOUND pipeline, shown at fixed time-points.

Furthermore, the addition of SOUND filtering generally resulted in greater retention of variance relative to the Original pipeline (Table 1,Tables S8, S9), and resulted in the removal of fewer ICs, particularly following the second round of ICA.

**Table 1:** Percentage variance removed and components removed (number of components / total components * 100) by each ICA step (ICA step 1, ICA step 2) in the Original and SOUND pipelines for each condition. Differences between pipelines shown in column Diff. * denotes significant difference between pipelines at p < 0.05 (Wilcoxon Signed Rank).

| Condition | Variance Removed (%) |  |  | Components Removed (%) |  |  |
| --- | --- | --- | --- | --- | --- | --- |
|  | Orig. | SOUND | Diff | Orig. | SOUND | Diff |
| CB-DC | 46.85, 38.63 | 45.20, 48.75 | 1.65, -10.12 | 11.69, 46.12 | 11.58, 26.58 | 0.11, 19.54* |
| CB-Elec | 37.63, 37.35 | 28.60, 11.85 | 9.02, 25.50 | 3.92, 42.86 | 4.84, 18.53 | -0.92*, 24.32* |
| CB-F8 | 41.93, 29.78 | 33.64, 23.69 | 8.30, 6.08 | 9.85, 46.80 | 9.05, 23.66 | 0.79, 23.13* |
| OP-DC | 57.07, 58.02 | 37.38, 29.67 | 19.69*, 28.35* | 12.00, 48.41 | 10.24, 23.09 | 1.76, 25.31* |
| OP-F8 | 66.66, 44.47 | 38.70, 29.42 | 27.96*, 15.05 | 7.50, 50.81 | 8.11, 28.67 | -0.61, 22.14* |

In summary, cerebellar stimulation with a DC coil results in considerable artefacts in the EEG trace. The addition of SOUND filtering to the cleaning process was necessary to improve performance, and thus only data from this pipeline were used in forthcoming analyses.

### Comparisons Across Conditions

#### TMS vs multisensory control

Figure 5 shows TEPs across all channels at different time points for all conditions. To estimate the extent of sensory-evoked activity following stimulation of the cerebellum and occipital/parietal region, responses following active stimulation were spatially and temporally correlated with responses following the multisensory control condition in sensor space. While the spatial distributions of evoked potentials appeared to differ between conditions at early latencies, similarities were evident across all conditions, including the multisensory control, from as early as 48 ms post-pulse (spatial correlations - Figure 6A). Analyses for each channel across time (temporal correlation) showed stronger relationships between TMS and the multisensory control condition in the mid (60-180 ms) and late (180-280 ms) latency periods relative to early (20-60 ms) latencies in all conditions (Figure 6B). At early latencies, the CB-DC condition showed strong correlations with the multisensory control at mid-central sites, and near the site of stimulation. Values were lowest at a contralateral posterior-parietal site (electrodes P1, P3, PO3) and a mid-ipsilateral site (electrodes FC6, C6).

**Figure 6:**
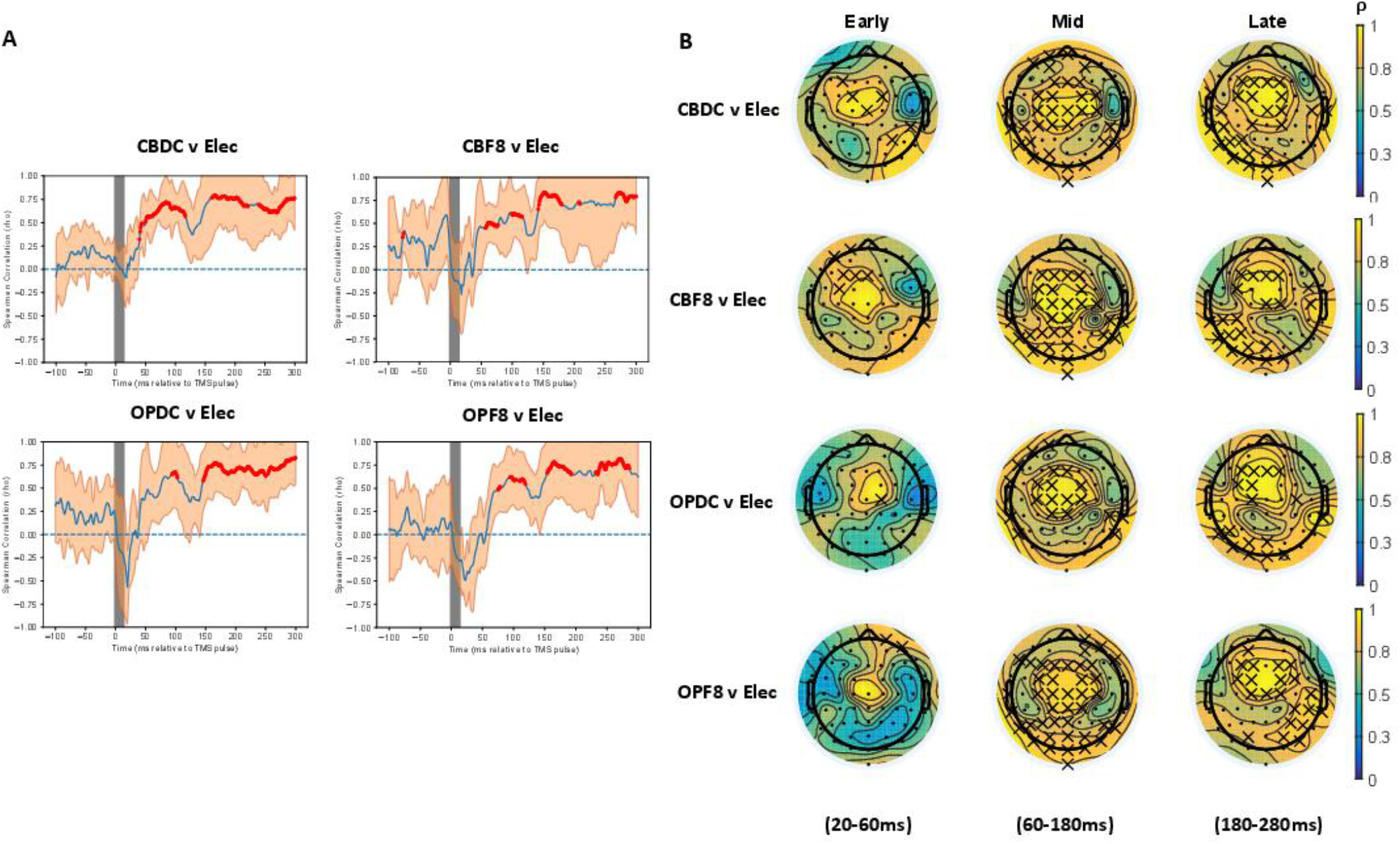
A. Spatial correlations (grand average) for all TMS conditions with electrical control following cleaning with the SOUND pipeline. Red dots indicate significant positive correlations. No negative correlations were found. B. Temporal correlations for each active TMS condition against the electrical control following cleaning with the SOUND pipeline. Crosses indicate significant positive correlations. No negative correlations were found.

Next, we assessed whether using a different referencing scheme impacted the correlations between the different conditions and the multisensory control. The surface Laplacian was chosen as it attenuates the effect of volume conduction at each electrode. Laplacian filtering only moderately reduced spatial correlations, but reduced the number of electrodes showing significant temporal correlations in the CB-DC condition, particularly at early latencies (Figure S5). Reductions were also observed at later latencies, but primarily over occipital electrodes.

Source estimation of the data confirmed that the spatial and temporal features of the TMS-evoked and multisensory control signals differed at early latencies, but were similar at later latencies (Figure 7A). However, early differences were less pronounced in the CB-F8 condition suggesting a dominance of sensory-evoked potentials in this condition (Figure 7B). Notably, all TMS conditions showed a greater response over occipital-parietal electrodes relative to the multisensory control at the latency corresponding to the first peak in the timeseries (Figure 8). The CB-DC condition evoked a higher degree of activity in bilateral frontal and left-hemispheric areas relative to the multisensory control, in conjunction with greater activity at the site of stimulation.

**Figure 7:**
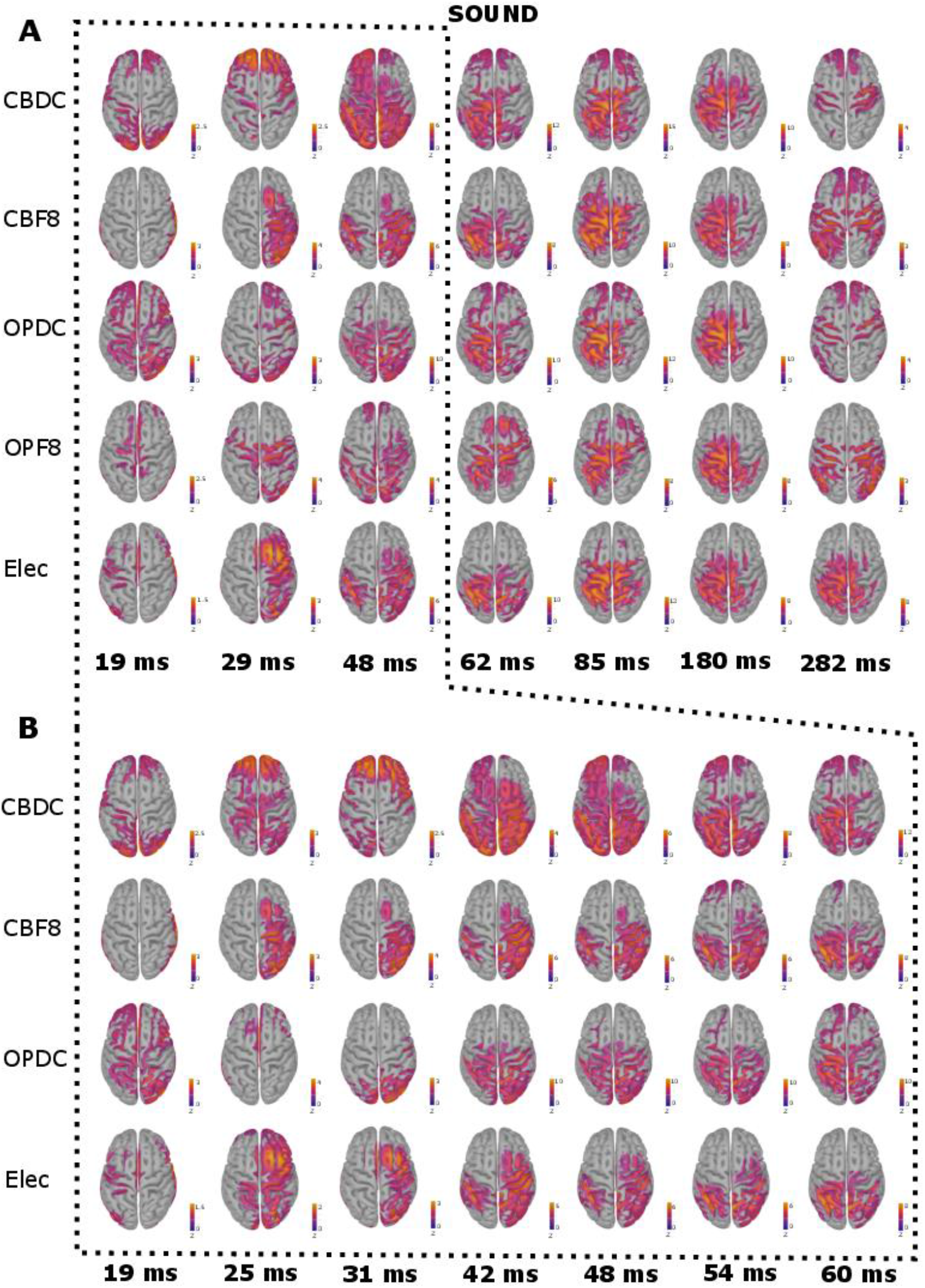
A. Estimated source distributions for the TMS and electrical control conditions using minimum norm estimation (MNE) following the SOUND pipeline, shown over full time-range. B. Early-latency estimated source distributions for the cerebellar TMS, Double-Cone TMS and electrical control conditions following the SOUND pipeline. MNE maps are thresholded to 40% of the maximum activity at each time point (minimum size of 50 vertices for activated region), and presented as z-scores. Colour scales show from 0 – maximum score for that time-point. Note the greater resemblance of the CB-F8 condition to the electrical control relative to the CB-DC condition.

**Figure 8:**
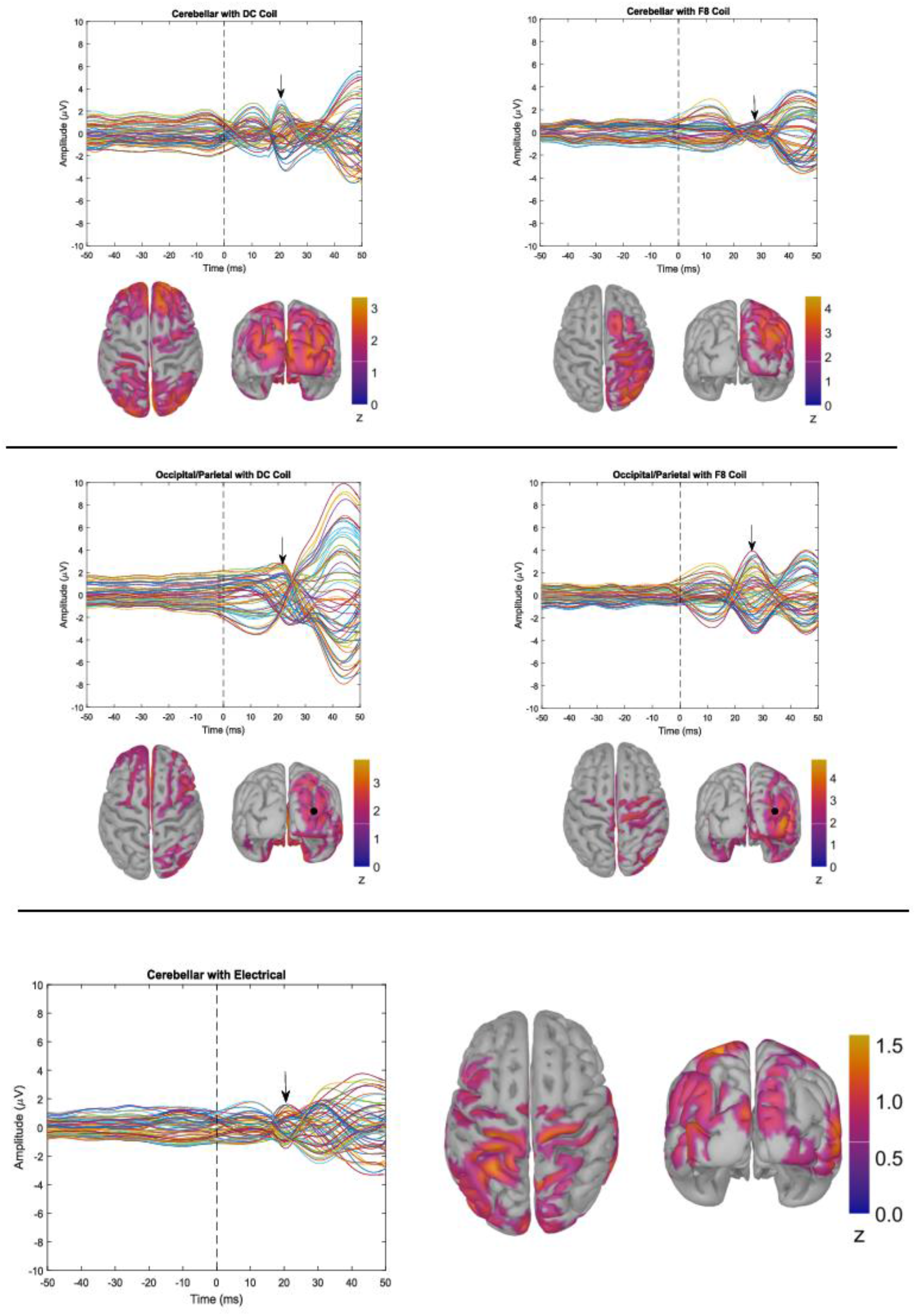
Source estimates corresponding to the first time-locked peak (from 15 ms post-pulse) in the timeseries for each condition (indicated by arrow). Colour scales show from 0 – maximum z-score for that time-point. Location of occipital/parietal stimulus shown as a black dot. The source distribution for TMS conditions revealed greater activity close to the site of stimulation relative to the electrical control, which elicited more activity in central regions, typical of a sensory response.

In summary, we found that all conditions showed high correlations with the multisensory control at mid to late latencies in both sensor and source space regardless of referencing montage. This suggests that latter time points largely reflected a generalised sensory response to stimulation. In contrast, correlations were lower and less widespread in the first 50 ms following stimulation, suggesting some difference between active and sensory stimulation protocols.

### Stimulation Site and Coil-Type

To determine if the response profile of the DC coil at the cerebellar stimulation site differed to those of the other active TMS conditions, comparisons were made within coil-type (across sites) and within stimulation site (across coil-type). Given the high correlations with the multisensory control for all conditions at later latencies, we focused on the first 60 ms following the TMS pulse. First, we assessed whether the responses following DC coil stimulation over the cerebellum (i.e., CB-DC) differed from the multisensory control condition, and DC control site (i.e., OP-DC). Cluster-based permutation tests on the Laplacian filtered data highlighted a significant difference between the CB-DC condition and multisensory control which was largest over posterior and midline channels (electrodes P1, P3, CP1) between 42-51 ms, extending to channels PO3 and CP3 between 51-60 ms (n = 9, p < 0.01) (Figure 9). Significant differences were also found between 51-60 ms over right parietal channels (electrodes PO4, PO8, P2, P4, CP2, CP4) (n = 9, p = 0.01). When comparing the CB-DC to the OP-DC conditions, differences were similarly observed over parietal electrodes (electrodes PO3, P1, P3, P5), although this did not reach the a priori threshold for statistical significance (n = 10, p = 0.27). Of note, no similar region was identified when comparing the OP-DC condition against the multisensory control (Figure S6). Source estimates for the CB-DC, and OP-DC conditions between 19-60 ms showed that the distribution of activity was generally more widespread following cerebellar stimulation at ~42 ms (Figure 7B), particularly in posterior and frontal regions. Taken together, these data suggest an early activation in left parietal electrodes that is unique to the CB-DC condition.

**Figure 9:**
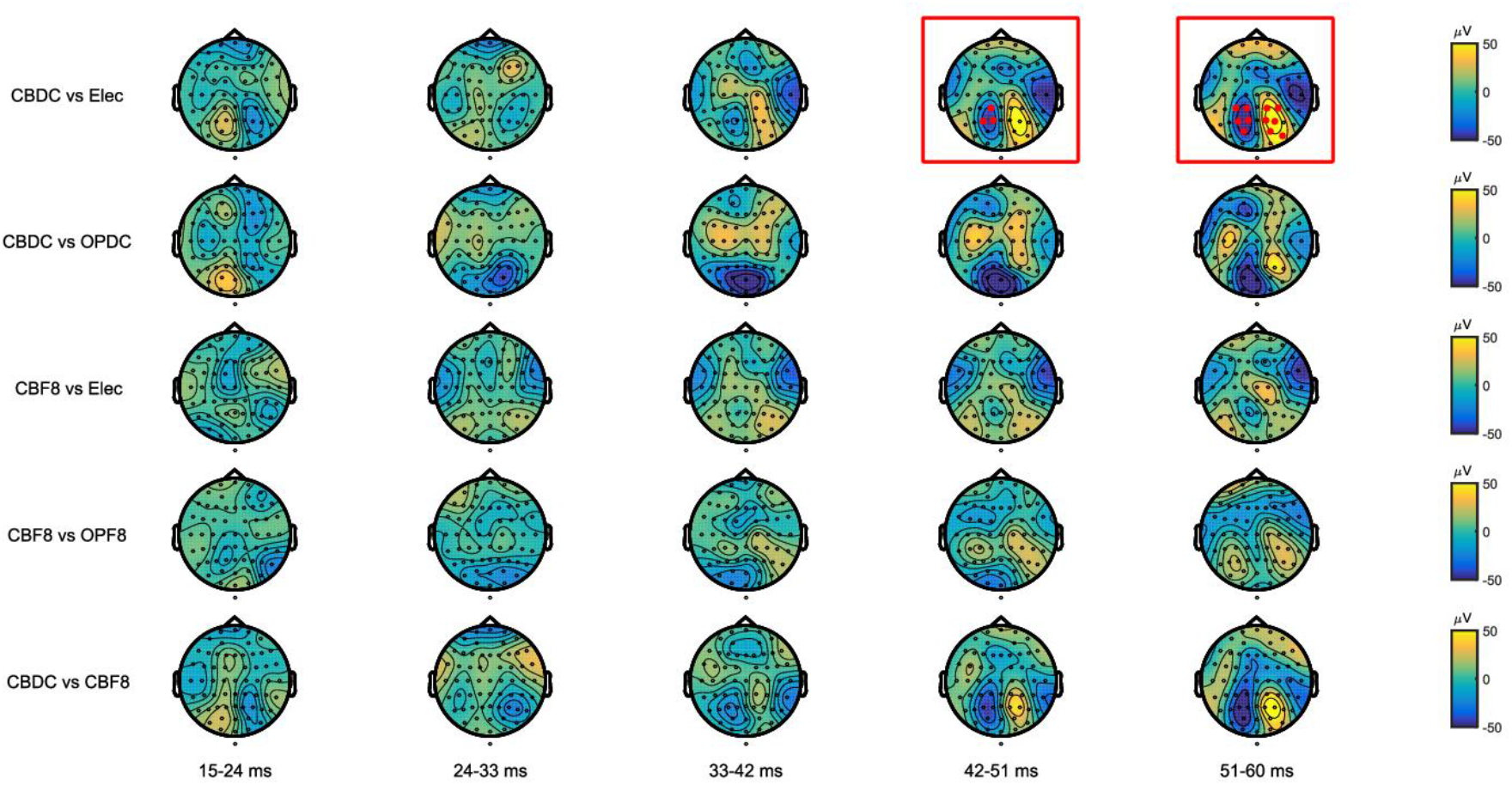
Cluster-based permutation tests comparing voltage distributions of peaks between 15-60 ms following cleaning with the SOUND pipeline and Laplacian filtering. Maps show voltage differences between conditions. Clusters defined as at least two adjacent electrodes exceeding p-value < 0.05 at each time-point. Critical alpha level for Monte-Carlo p-values set at p<0.025, 5000 iterations. Red dots indicate electrode clusters showing a statistically significant difference between conditions – these maps are highlighted by red squares. Time-windows were pre-defined.

Next, we assessed whether the response to F8 coil stimulation of the cerebellum differed between conditions. We could not find evidence for a difference between the F8 coil and the multisensory control at early latencies using cluster-based permutation tests (i.e., CB-F8 vs Elec, Figure 9). Comparisons across stimulation sites (i.e., CB-F8 vs OP-F8) also showed no evidence for a difference between conditions, suggestive of a common response profile for this coil across both cerebellar and parietal regions, which was largely dominated by sensory components.

Finally, we assessed whether responses to DC coil stimulation differed from the F8 coil at the cerebellar site. Cluster-based permutation tests on the Laplacian data taken across coil types (i.e., CB-DC vs CB-F8) highlighted a larger peak over parietal electrodes at latencies from 41-60 ms for the CB-DC condition which approached significance (n = 8, p = 0.03) (Figure 9). These results are consistent with a unique pattern following cerebellar stimulation with the DC coil.

### Suppression of Sensory Response

To further assess whether any TEPs were independent from sensory activity, SSP-SIR was used on the common referenced signals to suppress signals present in the electrical control condition from the CB-DC condition (Figures 10). Activity over occipital-parietal electrodes was heavily suppressed (Figure 10B). However, filtering did not substantially alter the temporal profiles of the CB-DC timeseries, whereby the most prominent peaks were generally preserved, albeit with a marked reduction in voltage amplitude (Figures 10A). Source estimates revealed focal activity in a contralateral parietal area at early latencies (Figure 10B), consistent with the parietal cluster identified by permutation analyses against the multisensory control (Figure 9).

**Figure 10:**
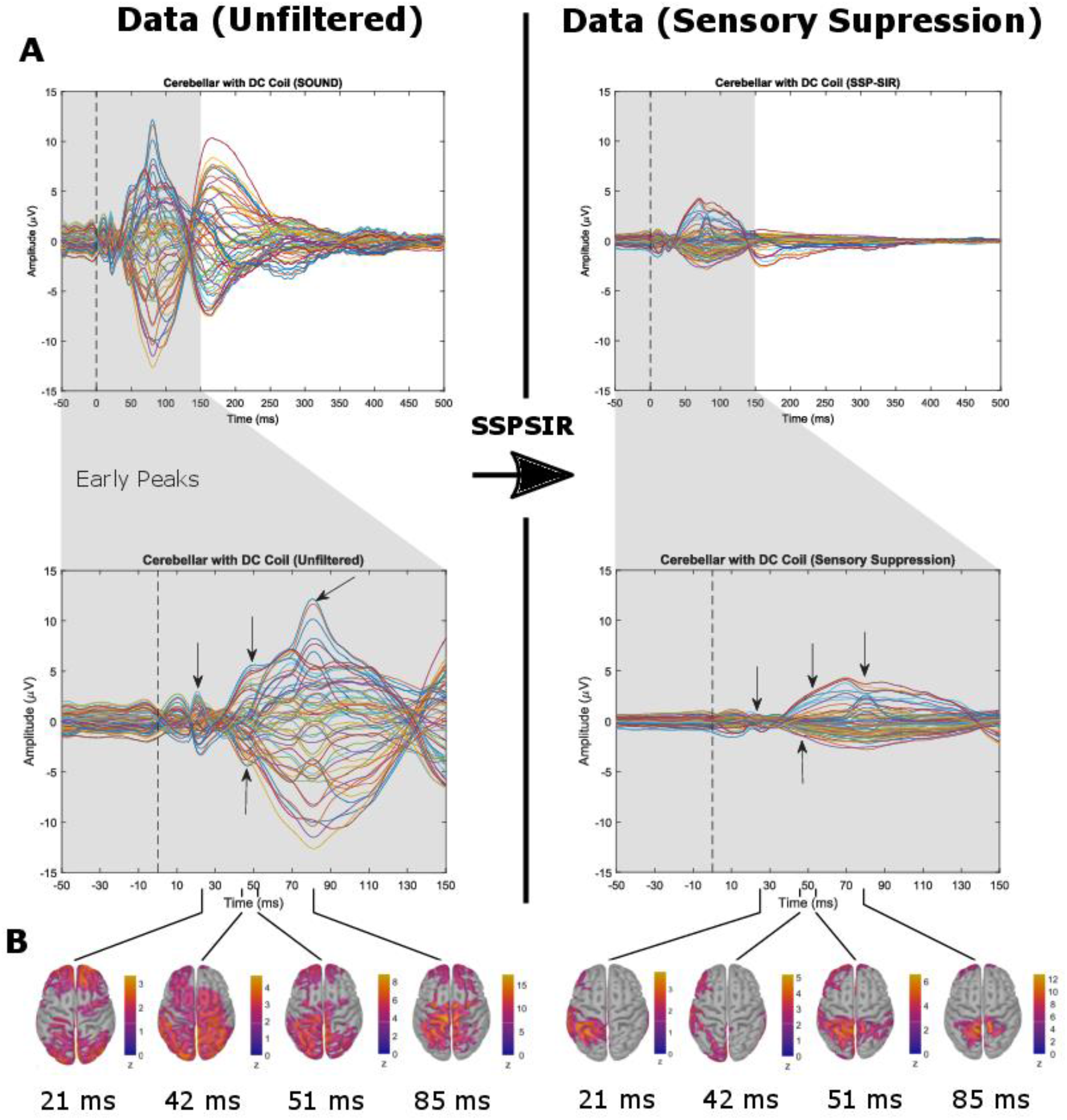
Peaks following SOUND and SSP-SIR filtering for the CB-DC condition. A. Timeseries at full range (−50 – 500 ms post-pulse) and magnified to show earlier peaks (−50 – 150 ms). Arrows highlight most prominent peaks. Note that peaks were retained, but attenuated following SSP-SIR. B. Source estimates corresponding to prominent early latency peaks for the CB-DC condition following the SOUND pipeline, and subsequent SSP-SIR filtering. Colour scales show from 0 – maximum z-score for that time-point.

## Discussion

This study assessed the feasibility of using TEPs to explore cerebellar-cortical connectivity via combined single-pulse cerebellar TMS using a DC coil with cortical EEG. We found that the DC coil produced large voltage deflections and offsets in the raw signal at both the cerebellar and an occipital/parietal control site. Using a F8 coil for the cerebellar stimulus generally reduced voltage amplitudes, and was deemed more comfortable by participants, but the high intensities necessary to ensure sufficient activation of cerebellar tissue resulted in a large portion of the EEG signal being dominated by sensory components. The addition of SOUND filtering to the cleaning pipeline reduced the signal offsets and resulted in more distinct peaks in the signals, suggesting that this pre-processing method is useful for highly artifactual data. Potentials from all active conditions were highly correlated with the multisensory condition after ~50 ms, suggesting a large amount of the late evoked response following cerebellar and cortical stimulation results from sensory input. However, weaker correlations were observed at earlier latencies, and Laplacian filtering revealed a parietal component unique to the CB-DC condition at ~42 ms. This was further evidenced by the suppression of the perceptual components via SSP-SIR on the SOUND cleaned (average referenced) data and source analysis, suggesting cerebellar stimulation with a DC coil may evoke activity in the parietal cortex. Generally, our findings suggest that cerebellar stimulation with a DC coil produces large artefacts in the EEG trace. While these artefacts can be suppressed to some extent by state-of-the-art cleaning methods, a large portion of the resultant components appear to be contaminated, possibly relating to the sensory and muscle-related response to the coil discharge, and spread to occipital regions. Whether ensuing components reflect long-range network activity arising from TMS activation of the cerebellarthalamocortical tract, or are dominated by sensory contamination and/or coil-driven artefacts (including incidental activation of additional proximal brain regions and associated network spread) remains open to further investigation.

Consistent with our present findings, stimulation with a DC coil at intensities required to reach cerebellar tissue is reportedly uncomfortable [8, 22]. Stimuli applied to the posterior fossa results in pronounced facial and neck muscle twitch, and often large eye-blink, while positioning the coil over cerebellar hemispheres stimulates lateral scalp muscles. In addition to TMS-evoked muscle activation, we observed participants producing voluntary movements in response to the stimuli, such as clenching a hand, or moving away from the coil. This likely had a detrimental effect on the integrity of the EEG signal due to associated muscle activity, and electrode displacement.

Voltage deflections provoked by the concurrent TMS stimuli were large in all conditions, and did not significantly differ between coil types at the cerebellar site. Whilst the largest, early peak (i.e., <10 ms) likely reflected TMS-evoked muscle activity, other muscle-related activity (such as jaw clenching, facial twitch), and eyeblinks also visibly contaminated the traces. We additionally found that the use of a DC coil increased the time taken for voltage offsets to return to baseline levels relative to the F8 coil.

Due to the considerable artefact profile of cerebellar TMS-EEG, the choice of artefact - suppression methods is a crucial factor for determining the feasibility of cerebellar-cortical assessment using this method. We found that the addition of SOUND filtering prior to ICA substantially improved the processed signal. As the SOUND filter increases the signal-to-noise ratio specific to individual electrodes, component estimation was likely aided by the reduction of decay artefacts and line noise within channels. However, in the absence of ground truth, we cannot assess how much of the underlying brain signal of interest was removed by these processes. As in previous studies [12, 14, 44], we found high spatiotemporal similarity between cleaned active TMS signals and the multisensory control at mid-to-late latencies, suggesting a common source within these time intervals. High correlations were observed primarily at the vertex, consistent with the N100 evoked potential. This component has been reported following active and sham stimulation, and is invariant to stimulus location (scalp or periphery) [12, 14, 44]. The spatiotemporal distributions of the electrical stimulus were also consistent with those reported following prefrontal and parietal stimulation in Conde, Tomasevic (12). The similarity between TMS and electrically-evoked distributions therefore suggests a common sensory source arising from the activation of scalp nerve fibres and auditory circuits.

Prior to 50 ms, we observed variations in spatiotemporal distributions across all conditions, and a parietal component was revealed that appeared to be specific to cerebellar stimulation with a DC coil. Structural and functional connectivity between the posterior cerebellum and posterior parietal cortex has been demonstrated through viral tracing methods in non-human primates [45] and human fMRI [46, 47]. One combined theta-burst stimulation (TBS) and EEG study found similar bidirectional effects on TEPs evoked from both the posterior parietal cortex and M1 following cerebellar cTBS and iTBS, suggesting that these regions are connected via a common modular circuit [48]. In the current study we did not observe distinct activity over motor regions following cerebellar DC stimulation, as might be expected. However, as 60 %MSO is a lower bound for TMS-induced cerebellar suppression of MEPs [22], it is possible that the stimulus delivered by the DC coil in the current study did not sufficiently activate Purkinje cells in anterior (motor) portions of the cerebellar cortex, as the DC coil sat several (~10) millimetres further away from the scalp than in typical CBI experiments due to both the addition of a foam covering over the face of the coil, and the EEG cap. Alternatively, studies on cerebellar suppression of MEPs have shown that cerebellar-to-M1 conduction times following an electromagnetic stimulus are in the order of 5-7 ms [7]. Therefore, the peak response in motor regions may not have been visible due to the large voltage deflections produced by the TMS pulse between 0-15 ms (where this interval was removed and interpolated in the cleaning process).

If not related to TMS activation of the cerebellarthalamocortical tract, the observed parietal component may have reflected a pulse-related artefact unique to the DC coil, or activation of the dorsal parietal stream from visual cortex stimulation spreading from the stimulation site due to the broad field of the coil. The similarities between the CB-DC and OP-DC conditions revealed by cluster-based permutation tests do indicate a degree of influence of coil-type on the evoked response. Regarding differences across coil-types for the cerebellar site, as both DC and F8 coils were discharged at intensities capable of reaching cerebellar tissue, we may have expected greater similarity between the CB-DC and CB-F8 conditions. However, it is likely that the induced fields differed substantially between coils, and thus stimulated distinct cerebellar structures. Additionally, as the CB-F8 condition showed a high degree of similarity to the multisensory control at all latencies, it is likely that resultant potentials from the F8 coil were largely dominated by sensory components.

Some insight into the degree of sensory contamination of the CB-DC signal may be gained from the SSP-SIR output. Notably, potentials at parietal-occipital electrodes were heavily attenuated, suggesting that a substantial portion of the signal at these sites contained peripherally-evoked potentials. However, as SSP-SIR is an aggressive artefact removal method, the attenuation may have been the result of over-correction. Thus, as the extent of sensory contamination of the signal is currently unknown, caution should be taken when interpreting the EEG signal following cerebellar stimulation, particularly when using a DC coil. Reducing the intensity of the stimulus may to some extent mitigate muscle activation, but further investigation into lower bounds for activating posterior cerebellar regions using a variety of coil types is necessary. It must also be pointed out that some remnants of the coil click could still be heard in all conditions, and were likely amplified by the proximity of the coil to the participant’s right ear. Although auditory artefacts were controlled for by the inclusion of white noise masking and coil discharge in all control conditions, we cannot rule out slight differences in auditory characteristics due to coil positioning and type. To determine the true nature of the components observed in the current study, methods with greater spatial precision, such as intracranial electroencephalography, and animal models, are required. Means of uncovering very early latency responses (< 15 ms) are also crucial to the development of cerebellar TMS-EEG research.

The current study had some limitations. Firstly, due to the use of a small sample, inferential statistics were underpowered, and grand averages may not generalise well to the broader population. However, the sample size was deemed sufficient for use in a feasibility study to demonstrate the degree of individual variability following cerebellar TMS-EEG, and to provide a preliminary assessment of the extent to which artefact may be confounding the signal of interest following cerebellar TMS with a DC coil. Secondly, the use of ICA may have introduced spurious features into the data; however, as a previous study demonstrated that ICA did not significantly alter correlations between signals [14], we chose to include it in both processing pipelines. It is also likely that extra-cranial muscle and intracranial sensory components that were not temporally independent of the TMS-related brain activity were not removed by the ICA. We attempted to mitigate the influence of sensory artefact in our analyses by the use of realistic and active control conditions. However, our multisensory control may not have optimally triggered all TMS-evoked sensory components at cerebellar and occipital/parietal sites due to differences in the level of activation of muscles and sensory pathways between TMS and electrical stimuli. While we may suggest using larger currents to elicit greater activation of neck and facial muscles, as stimuli were delivered at intensities within a tolerable range, increasing stimulus intensity may compromise ethical standards, and risk the direct activation of neural and/or spinal tissue. These points highlight a limitation of this form of somatosensory control more generally. Lastly, our source estimates should be interpreted cautiously, as we did not build individual structural MRI into the estimation model, but used a general template.

## Conclusion

This is the first study to systematically assess the feasibility of obtaining TEPs from cerebellar TMS-EEG using a realistic control condition, and state-of-the-art cleaning methods. Our results showed that cerebellar stimulation with a DC coil resulted in very large amplitude artefacts in the EEG trace. We found that the spatiotemporal distribution of responses in cerebral regions were highly correlated with the electrical control condition from ~50 ms post-pulse, suggesting a large contribution of sensory-evoked potentials to the overall signal after this time-point. However, components unique to the TMS condition were revealed prior to 50 ms, and were located over the parietal cortex. Whether these components largely reflected activation of the cerebellarthalamocortical tract, or were dominated by sensory contamination and/or coil-driven artefacts could not be reliably determined in the current study. As such, further research into the neurological and artefactual nature of these early components using additional technologies are necessary, in conjunction with methods to minimise the large muscle and sensory response evoked by the TMS pulse. Given the findings of the current study, we cannot recommend the use of single-pulse cerebellar TMS with EEG until the development of more sophisticated methods of artefact removal, or means of reducing sensory and muscle-related responses to coil discharge. Alternatively, ‘offline’ technologies, such as theta burst stimulation (TBS) may be considered, as these elicit persisting modulatory effects on neural excitability and thus may be applied prior to EEG recording to circumvent pulse-related artefacts inherent in TMS-EEG. Until such research has been conducted, all outcomes of cerebellar single-pulse TMS with cerebral cortical EEG should be treated with caution.

## Acknowledgements

The authors thank Ms Teresa Bollard and Ms Tabitha McNamara for help with data collection. N.C.R. is funded by a DECRA fellowship from the Australian Research Council (DE180100741). P.G.E. is funded by a Future Fellowship from the Australian Research Council (FT160100077).

## References

1. Van Overwalle F, Van de Steen F, Marien P. Dynamic causal modeling of the effective connectivity between the cerebrum and cerebellum in social mentalizing across five studies. Cogn Affect Behav Neurosci. 2019;19(1):211–23.

2. Van Overwalle F, De Coninck S, Heleven E, Perrotta G, Taib NOB, Manto M, et al. The role of the cerebellum in reconstructing social action sequences: a pilot study. Soc Cogn Affect Neurosci. 2019;14(5):549–58.

3. Heleven E, van Dun K, Van Overwalle F. The posterior Cerebellum is involved in constructing Social Action Sequences: An fMRI Study. Sci Rep. 2019;9(1):11110.

4. Dum RP, Strick PL. An unfolded map of the cerebellar dentate nucleus and its projections to the cerebral cortex. Journal of Neurophysiology. 2003;89:634–9.

5. Stoodley CJ. The cerebellum and cognition: evidence from functional imaging studies. Cerebellum. 2012;11(2):352–65.

6. Stoodley CJ, Valera EM, Schmahmann JD. Functional topography of the cerebellum for motor and cognitive tasks: an fMRI study. Neuroimage. 2012;59(2):1560–70.

7. Ugawa Y, Yoshikazu U, Terao Y, Hanajima R, Kanazawa I. Magnetic stimulation over the cerebellum in humans. Annals of Neurology. 1995;37(6):703–13.

8. Hardwick RM, Lesage E, Miall RC. Cerebellar transcranial magnetic stimulation: the role of coil geometry and tissue depth. Brain Stimul. 2014;7(5):643–9.

9. Fernandez L, Major BP, Teo WP, Byrne LK, Enticott PG. Assessing cerebellar brain inhibition (CBI) via transcranial magnetic stimulation (TMS): A systematic review. Neurosci Biobehav Rev. 2018;86:176–206.

10. Chung SW, Rogasch NC, Hoy KE, Fitzgerald PB. Measuring Brain Stimulation Induced Changes in Cortical Properties Using TMS-EEG. Brain Stimul. 2015;8(6):1010–20.

11. Gordon PC, Desideri D, Belardinelli P, Zrenner C, Ziemann U. Comparison of cortical EEG responses to realistic sham versus real TMS of human motor cortex. Brain Stimul. 2018;11(6):1322–30.

12. Conde V, Tomasevic L, Akopian I, Stanek K, Saturnino GB, Thielscher A, et al. The non-transcranial TMS-evoked potential is an inherent source of ambiguity in TMS-EEG studies. Neuroimage. 2018;185:300–12.

13. Ilmoniemi RJ, Kicic D. Methodology for combined TMS and EEG. Brain Topogr. 2010;22(4):233–48.

14. Biabani M, Fornito A, Mutanen TP, Morrow J, Rogasch NC. Characterizing and minimizing the contribution of sensory inputs to TMS-evoked potentials. Brain Stimul. 2019;12(6):1537–52.

15. Rogasch NC, Fitzgerald PB. Assessing cortical network properties using TMS-EEG. Hum Brain Mapp. 2013;34(7):1652–69.

16. Rogasch NC, Sullivan C, Thomson RH, Rose NS, Bailey NW, Fitzgerald PB, et al. Analysing concurrent transcranial magnetic stimulation and electroencephalographic data: A review and introduction to the open-source TESA software. Neuroimage. 2017;147:934–51.

17. Fernandez L, Rogasch NC, Do M, Clark G, Major BP, Teo WP, et al. Cerebral Cortical Activity Following Non-invasive Cerebellar Stimulation-a Systematic Review of Combined TMS and EEG Studies. Cerebellum. 2020;19(2):309–35.

18. Rogasch NC, Thomson RH, Daskalakis ZJ, Fitzgerald PB. Short-latency artifacts associated with concurrent TMS-EEG. Brain Stimul. 2013;6(6):868–76.

19. Rogasch NC, Thomson RH, Farzan F, Fitzgibbon BM, Bailey NW, Hernandez-Pavon JC, et al. Removing artefacts from TMS-EEG recordings using independent component analysis: importance for assessing prefrontal and motor cortex network properties. Neuroimage. 2014;101:425–39.

20. Mutanen T, Maki H, Ilmoniemi RJ. The effect of stimulus parameters on TMS-EEG muscle artifacts. Brain Stimul. 2013;6(3):371–6.

21. Mutanen TP, Kukkonen M, Nieminen JO, Stenroos M, Sarvas J, Ilmoniemi RJ. Recovering TMS-evoked EEG responses masked by muscle artifacts. Neuroimage. 2016;139:157–66.

22. Fernandez L, Major BP, Teo WP, Byrne LK, Enticott PG. The Impact of Stimulation Intensity and Coil Type on Reliability and Tolerability of Cerebellar Brain Inhibition (CBI) via Dual-Coil TMS. Cerebellum. 2018.

23. Werhahn KJ, Taylor J, Ridding M, Meyer BU, Rothwell JC. Effect of transcranial magnetic stimulation over the cerebellum on the excitability of human motor cortex. Electroencephalography and Clinical Neurophysiology - Electromyography and Motor Control. 1996;101(1):58–66.

24. Oldfield R. The assessment and analysis of handedness: The Edinburgh inventory. Neuropsychologia. 1971;9(1):97–113.

25. Rossi S, Hallett M, Rossini PM, Pascual-Leone A, Safety of TMSCG. Safety, ethical considerations, and application guidelines for the use of transcranial magnetic stimulation in clinical practice and research. Clin Neurophysiol. 2009;120(12):2008–39.

26. Taylor PC, Walsh V, Eimer M. The neural signature of phosphene perception. Hum Brain Mapp. 2010;31(9):1408–17.

27. Tiitinen H, Virtanen J, Ilmoniemi R, Kamppuri J, Ollikainen M, Ruohonen J, et al. Separation of contamination caused by coil clicks from responses elicited by transcranial magnetic stimulation. Clinical Neurophysiology. 1999;110(5):982–5.

28. Deng ZD, Lisanby SH, Peterchev AV. Electric field depth-focality tradeoff in transcranial magnetic stimulation: simulation comparison of 50 coil designs. Brain Stimul. 2013;6(1):1–13.

29. Çan MK, Laakso I, Nieminen JO, Murakami T, Ugawa Y. Coil model comparison for cerebellar transcranial magnetic stimulation. Biomedical Physics & Engineering Express. 2018;5(1).

30. Ugawa Y, Day BL, Rothwell JC, Thomson PD, Merton PA, Marsden CD. Modulation of motor cortical excitability by electrical stimulation over the cerebellum in man. Journal of Physiology. 1991;441:57–72.

31. Delorme A, Makeig S. EEGLab: an open source toolbox for analysis of single-trial EEG dynamics including independent component analysis. Journal of neuroscience methods. 2004;134(1):9–21.

32. Mutanen TP, Biabani M, Sarvas J, Ilmoniemi RJ, Rogasch NC. Source-based artifact-rejection techniques available in TESA, an open-source TMS-EEG toolbox. Brain Stimul. 2020;13(5):1349–51.

33. Mutanen TP, Metsomaa J, Liljander S, Ilmoniemi RJ. Automatic and robust noise suppression in EEG and MEG: The SOUND algorithm. Neuroimage. 2018;166:135–51.

34. [dataset] Fernandez L. Assessing cerebellar-cortical connectivity using concurrent TMS-EEG: A Feasibility Study, 2020. 10.6084/m9.figshare.13082495.

35. Tadel F, Bock E, Niso G, Mosher J, Cousineau M, Pantazis D, et al. MEG/EEG Group Analysis with Brainstorm. Frontiers in Neuroscience. 2019.

36. Dunn OJ, Clark V. Correlation coefficients measured on the same individuals. Journal of the American Statistical Association. 1969;64(325):366–77.

37. Charter RA, Larsen BS. Fisher's z to r. Educational and Psychological Measurement. 1983;43(1):41–2.

38. Blair R, Karniski W. An alternative method for significance testing of waveform difference potentials. Psychophysiology. 1993;30:518–24.

39. Oostenveld R, Fries P, Maris E, Schoffelen J. FieldTrip: open source software for advanced analysis of MEG, EEG and invasive electrophysiological data. Comput Intell Neuroscience. 2011;2011(156869).

40. Cohen MX. Comparison of different spatial transformations applied to EEG data: A case study of error processing. Int J Psychophysiol. 2015;97(3):245–57.

41. Perrin F, Pernier J, Bertrand O, Echallier J. Spherical splines for scalp potential and current density mapping. Electroencephalography and Clinical Neurophysiology. 1989;72:184–7.

42. Winkler I, Debener S, Muller K-S, Tangermann M. On the influence of high-pass filtering on ICA-based artifact reduction in EEG-ERP. 37th Annual International Conference of the IEEE Engineering in Medicine and Biology Society (EMBC): IEEE; 2015.

43. Metsomaa J, Sarvas J, Ilmoniemi RJ. Multi-trial evoked EEG and independent component analysis. J Neurosci Methods. 2014;228:15–26.

44. Du X, Choa FS, Summerfelt A, Rowland LM, Chiappelli J, Kochunov P, et al. N100 as a generic cortical electrophysiological marker based on decomposition of TMS-evoked potentials across five anatomic locations. Experimental Brain Research. 2017;235(1):69–81.

45. Clower DM, Dum RP, Strick PL. Basal ganglia and cerebellar inputs to ‘AIP’. Cereb Cortex. 2005;15(7):913–20.

46. O'Reilly JX, Beckmann CF, Tomassini V, Ramnani N, Johansen-Berg H. Distinct and overlapping functional zones in the cerebellum defined by resting state functional connectivity. Cereb Cortex. 2010;20(4):953–65.

47. Buckner RL, Krienen FM, Castellanos A, Diaz JC, Yeo BT. The organization of the human cerebellum estimated by intrinsic functional connectivity. J Neurophysiol. 2011;106(5):2322–45.

48. Casula EP, Pellicciari MC, Ponzo V, Stampanoni Bassi M, Veniero D, Caltagirone C, et al. Cerebellar theta burst stimulation modulates the neural activity of interconnected parietal and motor areas. Scientific Reports. 2016;6.

