## Supplementary Material for "Assessing cerebellar-cortical connectivity using concurrent TMS-EEG: A Feasibility Study"

***1. Supplementary Material – Original Pipeline:***

1) Remove unused electrodes

2) Save original channel locations

3) Remove bad electrodes (joint probability, threshold=5, measure=kurtosis, norm=on)

4) Epoching (-1000 ms to 1000 ms around pulse)

5) Remove baseline [-500 -10ms] relative to TMS pulse

6) Data removed [-2 10ms] (TMS-related muscle artefact), and interpolated using cubic spline fitting

7) Data downsampled to 1,000 Hz

8) Remove bad trials (joint probability and manual inspection)

9) ICA with component selection for TMS-evoked muscle activity (interpolated data section replaced with constant amplitude)

10) Data removal extended to [-2, 15ms] for additional artefacts, interpolated

11) Filter data (zero-phase Butterworth): Bandpass (1-100 Hz), bandstop (48-52 Hz)

12) ICA with component selection for remaining artefacts (interpolated data section replaced with constant amplitude)

13) Interpolate missing channels

14) Re-reference to common average reference

*Supplementary Material - Pipeline With SOUND Filtering:*

1) Remove unused electrodes

2) Save original channel locations

3) Epoching (-1000 ms to 1000 ms around pulse)

4) Remove baseline [-500 -10ms] relative to TMS pulse

5) Data removed [-2 10ms] (TMS-related muscle artefact), and interpolated using cubic spline fitting

6) Data downsampled to 1,000 Hz

7) SOUND filter (lambda = 0.1, iterations = 10)

8) Remove bad electrodes (joint probability, threshold=5, measure=kurtosis, norm=on)

9) Remove bad trials (joint probability and manual inspection)

10) ICA with component selection for TMS-evoked muscle activity (interpolated data section replaced with constant amplitude)

11) Data removal extended to [-2, 15ms] for additional artefacts, interpolated

12) Filter data (zero-phase Butterworth): Bandpass (1-100 Hz), bandstop (48-52 Hz)

13) ICA with component selection for remaining artefacts (interpolated data section replaced with constant amplitude)

14) Interpolate missing channels

15) Re-reference to common average reference

#### *2. Supplementary Material - Motor Response (1 mV)*

Resting motor response was assessed prior to the TMS-EEG trials via the first dorsal interosseous (FDI) muscle of the right hand (i.e., left M1) using a tendon-belly montage with the ground electrode placed over the styloid process of the ulna. Electromyographic (EMG) activity was recorded with a PowerLab data acquisition unit (ADInstruments, Dunedin, NZ) and LabChart8 software (ADInstruments, Dunedin, NZ), amplified x1000, filtered (lowpass = 0.3 Hz, highpass = 1 kHz), and digitised (10 kHz). TMS pulses were delivered to the left M1 via a Magstim 70 mm F8 coil (Ref. 3190-00). Trials were epoched -200 ms to 500 ms relative to each pulse. Threshold was defined as the minimum stimulation intensity necessary to induce an average MEP peak-to-peak amplitude of 1 mV over 10 trials. Electromyographic activity continued to be monitored throughout the session.

#### *3. Supplementary Material – Procedure for Determining Occipital/Parietal Site*

To ensure no phosphenes or other visual disturbances occurred when stimulating the occipital/parietal region, a standard procedure was followed in each session to determine the stimulation site used in a session. Specifically, all overhead lights in the room were turned off, and the participant was instructed to focus on the image of a white cross overlayed on a black background and placed at eye-level, approximately 50 cm from the seated individual. An initial stimulation location at 10-20 EEG site PO4 was chosen, and five pulses were delivered at trial intensity. Notably, the DC coil was rotated ~45 degrees in a clockwise direction to avoid the right-most wing activating the M1 when positioned at this site. The participant was then asked if they had noticed any visual disturbances. If none were reported, the lights were turned on, and trials commenced at the location. Otherwise, the coil was moved upward in 1 cm increments until no visual disturbances was reported.

#### *4. Supplementary Material – Raw Data*

Two datasets were removed due to poor signal quality (the CB-F8 condition for participants 5 and 6). The remaining raw traces showed similar artefact profiles across all stimulation conditions. An early-latency voltage spike (~0.25 ms post-pulse) was followed by a large amplitude peak occurring ~0.5 ms after pulse onset (Figure 3). The amplitude of this secondary spike was generally greatest in channels closest to the site of stimulation, decreasing in contralateral fronto-lateral regions. The voltage range of the artefact in the raw traces varied according to stimulation type, with the DC coil generally producing the largest deflections (Table S3). The amplitudes varied considerably across participants. Wilcoxon signed rank tests on voltage ranges (max-min) confirmed that the DC coil produced large deflections regardless of site (Table S4,S5: CB-DC vs OP-DC), and that ranges did not differ at the cerebellar site according to coil type (Table S4,S5: CB-DC vs CB-F8). All conditions except the OP-F8 produced significantly larger voltage deflections than the electrical control (Table S4,S5).

The initial large voltage peak was followed by a steep decline, and gradual recovery back to baseline levels (< ±150 µV). The range of recovery times [min, max] ms post-pulse varied substantially across conditions, CB-DC [1.5, 938.9] ms, CB-elec [0.8, 25.4] ms, CB-F8 [2.2, 108.3] ms, OP-DC [0.8, 207.0] ms, and OP-F8 [0.2, 913.9] ms. In rare instances, some channels failed to drop below 150 µV. This occurred primarily in the CB-DC condition and was likely a result of temporary poor skin-electrode contact due to movement. These channels were removed, and interpolated prior to further analysis. Wilcoxon signed rank tests confirmed that the DC coil produced larger recovery times than the F8 coil at the cerebellar site (Table S4,S5: CB-DC vs CB-F8). The DC coil also elicited greater recovery times at the cerebellar verses the occipital/parietal site (Table S4,S5: CB-DC vs OP-DC), where this was not the case for the F8 coil (Table S4,S5: CB-F8 vs OP-F8).

Blind source separation of the artefact via ICA generated independent components consistent with TMS-related muscle activation. This is also consistent with a large reduction in artefact during trials for a participant with a low tolerance to the electrical stimulation, whereby the low current (9.0 mA) failed to produce a sufficient twitch response (trials omitted from analysis). The large peaks were also absent in electrical stimulation trials when the DS7A stimulator failed to deliver a pulse due to poor electrode-to-skin contact (although the TMS coil still fired).

#### *5. Supplementary Material – Cleaning Pipelines*

Using a regularisation parameter of 0.1, the SOUND algorithm converged (relative change in sigma < 1%) within 10 iterations for 40 of the 47 datasets (85%). For the remaining datasets, SOUND was run using up to 20 iterations. Convergence was achieved in 11 iterations (one dataset), 12 (three), 13 (two), and 20 (one) respectively (though speed of convergence will depend on the initial estimate of the noise covariance matrix).

The proportion (of total) of trials and channels rejected for each condition following each pipeline is shown in Table S6. There was considerable variation in inter-subject signal profiles following both pipelines for all conditions. Supplementary Figure S3 shows the individual timeseries response following the Original pipeline, while Figure S4 shows individual responses following the SOUND pipeline. Looking at the grand average data (Figures 4, 5), distinct peaks at early latencies, and at ~100 ms and 200 ms were apparent following both pipelines. However, the maximum voltage ranges differed, whereby SOUND filtering consistently reduced the magnitude of extrema relative to the Original data cleaning sequence. The maxima and minima produced by each pipeline is shown in Table S7. SOUND filtering also reduced signal noise in the pre-stimulation and early (< 60 ms) periods (resulting in less ‘ringing’ artefact from bandpass filtering), and generally produced clearer peaks relative to the Original pipeline. This is particularly apparent in the active cerebellar conditions, and is also reflected in the spatio-temporal voltage distributions for each condition. Namely, SOUND filtering resulted in more distinct regions of activity at early latencies, and clearer highlighting of the response in the stimulated area.

Furthermore, the addition of the SOUND filtering process generally resulted in greater retention of variance relative to the Original pipeline (Table 1, Supplementary Table

S8, S9), although this was only significant for the OP-DC condition (ICA stages 1, 2), and the OP-F8 condition (ICA stage 1). Similarly, SOUND filtering generally resulted in the removal of fewer ICs relative to the Original pipeline, but this was predominantly significant following the second round of ICA.

### *6. Supplementary Material – Comparisons Across Conditions*

#### *6.1 TMS vs multisensory control*

To estimate the extent of sensory contamination remaining in traces from stimulation of the cerebellum with the DC coil following the SOUND pipeline, statistical comparisons with the electrical control were performed in sensor space. Correlation analyses for the TMS conditions versus multisensory control taken across spatial maps for each time-point (spatial correlation) showed positive values in pre-stimulus periods, followed by a sharp drop immediately following the TMS trigger and steep increase at around 50 ms (Figure 6A). Post 50 ms, relationships remained exceedingly positive (confidence intervals > 0) suggesting that the response to stimulation from both TMS coils and electrical current arose from a common source from this time-point.

Analyses confirmed that the potentials evoked by TMS were similar to those evoked by the electrical control across a 0-300 ms interval. When assessing mean differences at the cluster level, permutation tests on the SOUND data suggested that peaks differed primarily at mid to late latencies. Contrasts on both DC conditions (CB-DC and OP-DC) revealed that signals from the DC coil were significantly larger in amplitude than those from the control only after ~135 ms. From this time-point, the CB-DC condition showed the greatest dissimilarity to the electrical control in an ipsilateral (to the pulse) frontal region, before shifting to an area close to the site of stimulation and fronto-central regions, then moving to a contralateral site. Conversely, differences between the OP-DC condition and electrical control were initially most pronounced in a contralateral-central region before spreading centrally and toward the site of stimulation.

Laplacian filtering highlighted an additional negative cluster from ~42 ms when contrasting the CB-DC condition against the electrical control. This appeared as the leftmost side of a dipole at parietal electrodes (PO3, P1, P3, CP1, CP3), and suggests differences in parietal activity at the level of the dura. No other significant early-latency effects were found in the remaining conditions following Laplacian filtering.

Correlation analyses for the TMS versus control condition taken across spatial maps for each time-point (spatial correlation) generally showed positive correlations in pre-stimulus periods, followed by a sharp drop immediately following the TMS trigger and steep increase at around 50 ms (Figure 6A). Post 50 ms, relationships remained exceedingly positive (confidence intervals > 0) suggesting that the response to stimulation from both TMS coils and electrical current arose from a common source from this time-point. Laplacian filtering only marginally reduced early-latency correlations in all conditions (Figure S5).

Correlation analysis for each channel across time (temporal) similarly showed substantially higher significant relationships between DC and electrical conditions in the mid (60-180 ms) and late (180-280 ms) latency periods relative to early (20-60 ms) latencies in all conditions (Figure 6B). Values were predominantly highest over central and lateral channels, likely reflective of a common sensory response to the stimuli. At early latencies, the CB-DC condition showed high significant correlations with the electrical control at mid-central sites, and near the site of stimulation. Conversely, values were lowest in a mid-ipsilateral region, and around a contralateral parietal site identified in the permutation tests following Laplacian filtering. Early-latency correlation values taken between the OP-DC and electrical control were highest over a central site, and lowest at two mid-lateral regions (one contra-, one ipsilateral to the pulse), and close to the site of stimulation. Generally, values were considerably lower relative to the cerebellar site, likely because the control was optimised to mimic pulse effects to lateral cerebellar, rather than occipital site. Laplacian filtering reduced the number of significant temporal correlations in the cerebellar condition, particularly at early latencies, and reduced values at the contralateral parietal and right mid-lateral regions (Figure S5).

Source estimation of the SOUND data confirmed that the spatial and temporal features of both the TMS-evoked and electrical control signals became similar after approximately 85 ms (Figure 7A). However, conditions differed at early latencies (Figure 7B). The CB-DC condition evoked a higher degree of activity in bilateral frontal and left-hemispheric areas relative to the electrical control, in conjunction with greater activity at the site of stimulation.

#### *7. Supplementary Material – Stimulation Site and Coil-Type*

To determine if the response profile of the DC coil at the cerebellar stimulation site differed significantly to those of the other active TMS conditions, comparisons were made within coil-type (across sites) and within stimulation site (across coil-type). We found that the spatial and temporal profiles of signals evoked by respective coil types were similar at cerebellar and occipital/parietal stimulation sites. This was particularly apparent for the spatial distributions of signals evoked by stimuli delivered to the occipital/parietal region, where cross-coil correlation values remained positive at all post-pulse latencies (OP-DC verses OP-F8). Temporal correlation analyses confirmed this relationship, where a number of significant correlations were revealed at early latencies, predominantly in central and contralateral channels. While potentials evoked by cerebellar stimulation did not show the same degree of similarity at early latencies, both spatial and temporal correlation analyses revealed substantial similarities between coil types at mid to late latencies. These occurred predominantly at central and posterior sites in the temporal domain. Permutation testing showed that the DC coil produced a significantly larger signal in fronto-central cortical regions 150 ms post-pulse relative to the F8 coil following stimuli applied to either site, suggesting a greater response from this coil regardless of stimulation site.

Within coil (across site) correlations taken in the spatial domain generally displayed profiles resembling those of the TMS verses electrical control conditions, where values tended to increase post 50 ms. This was reflected in the temporal domain, whereby signals evoked by the DC coil showed significant correlations across sites only at mid and late-latencies in contralateral and posterior channels. The F8 coil similarly produced significant cross-site correlations at mid and late-latencies, predominantly in central, contralateral and posterior channels. Interestingly, permutation tests revealed that while potentials evoked 135-270 ms post-pulse by the DC coil were significantly higher when stimuli were applied to the cerebellar relative to the occipital/parietal site, there were no significant differences across sites for the F8 coil, suggestive of a common response profile for this coil.

Laplacian filtering did not substantially alter the trend of the spatial correlations across time in any comparison, but tended to produce more consistent values post 50 ms in the CB-DC verses CB-F8 and OP-DC verses OP-F8 comparisons ($\rho$ close to 0.5). Temporal correlations showed fewer significant values at late latencies due to removal of low-frequency components. Cluster permutation analyses did not reveal any additional early-latency spatiotemporal differences between conditions.

As sensor-level analyses suggested the DC coil may have elicited a distinct set of early-latency components when applied to the cerebellar site, we visually examined source estimates for the CB-DC, CB-F8, and OP-DC conditions between 19-60 ms (Figure 7). Comparing source distributions of the CB-DC condition with those of both the CB-F8 and OP-DC conditions indicated some coil-driven effects. Specifically, source distributions for the CB-F8 condition bore greater resemblance to those evoked by the electrical stimulus than the CB-DC condition prior to 50 ms, suggesting a large proportion of the early response evoked by this coil at the cerebellum may have been sensory, and not related to the response profile of the DC coil at this site. Conversely, the source distribution of the DC coil at the occipital/parietal site showed a similar profile to the cerebellar site at 19 ms. The TMS conditions showed similar responses after ~50 ms, consistent with correlation analyses.

### *8. Supplementary Material – Suppression of Sensory Response*

Correlations taken across site and coil-type in sensor space showed that SSP-SIR reduced signal similarity in the temporal domain across early, mid and late time periods for all comparisons. When comparing across stimulation sites for the DC coil at early latencies, high, but non-significant correlations were recorded at a left mid-parietal site, prefrontal and right temporal channels. The lowest values were recorded in contralateral parietal/occipital and frontal channels.

### *9. Supplementary Material - Source Estimation*

#### *9.1 DSPM – Not used in final study, but output available on request*

To form the source estimates for the cleaned and SSP-SIR data, a common head model was computed using a three-layer symmetric Boundary Element Method (BEM, OpenMEEG) using the default anatomy (ICBM152) and 1922 vertices per layer. Conductivities were set to the default values (scalp = 1, skull = 0.0125, brain = 1). Each individual’s datasets were then imported into Brainstorm, and averaged over epochs to obtain the time-locked traces for each condition. Noise covariance matrices were derived from the pre-stimulus recordings (-1s to -0.001s). Cortical sources were estimated using minimum norm estimation with dipoles assumed to be tangential to the surface. Amplitudes were normalised using dSPM resulting in unitless maps. These maps were then rectified to account for subject-specific amplitude and sign ambiguities resulting from the complex folding of the cortex. A weighted average was then performed over participants to form the grand average of a condition. As the averaging of N normalised maps reduces the variance by 1/N, the resultant averaged map was scaled to maintain a variance of 1 under the null hypothesis. This was used for display purposes only. Source estimates were smoothed with a kernel of size 3mm.

#### *10. Supplementary Material – Additional Images*

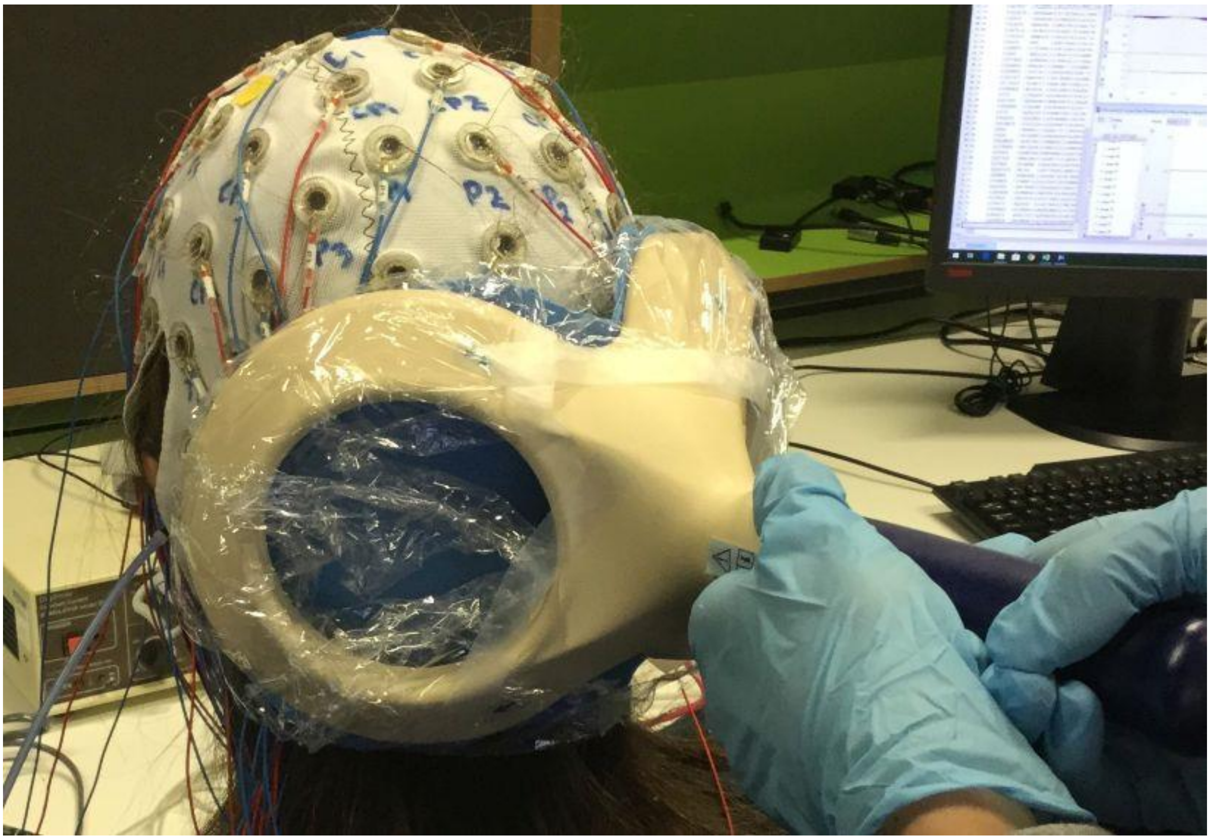

*Figure S1:* Double-cone TMS coil for cerebellar stimulation with concurrent EEG. A foam covering was placed over the surface of the coil to reduce to minimise contact with EEG electrodes, and dampen vibrations during coil discharge.

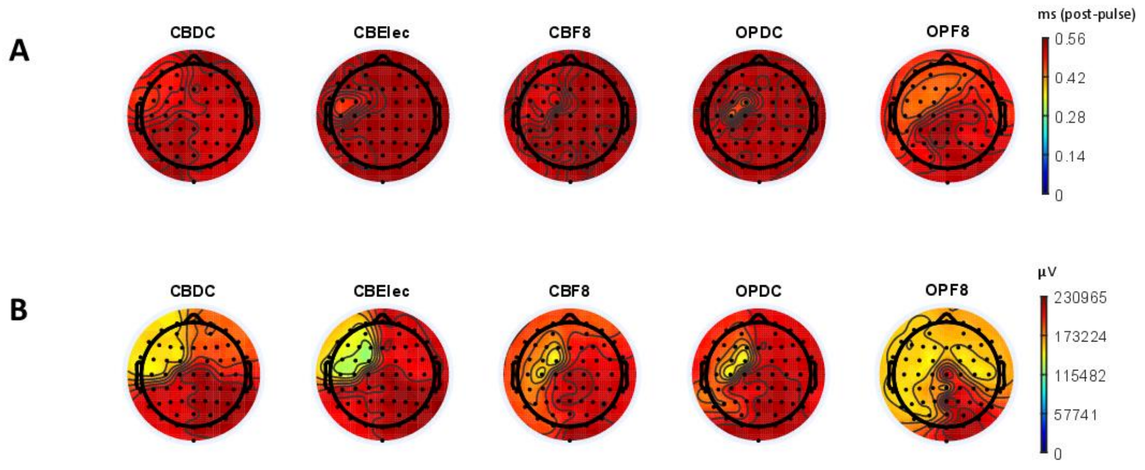


*Figure S2:* A. Times (ms post-pulse) for each channel (grand average) to reach maximum voltage in each condition. This occurs at approximately 0.5 ms in the majority of channels. B. Voltage maxima in each channel (µV).

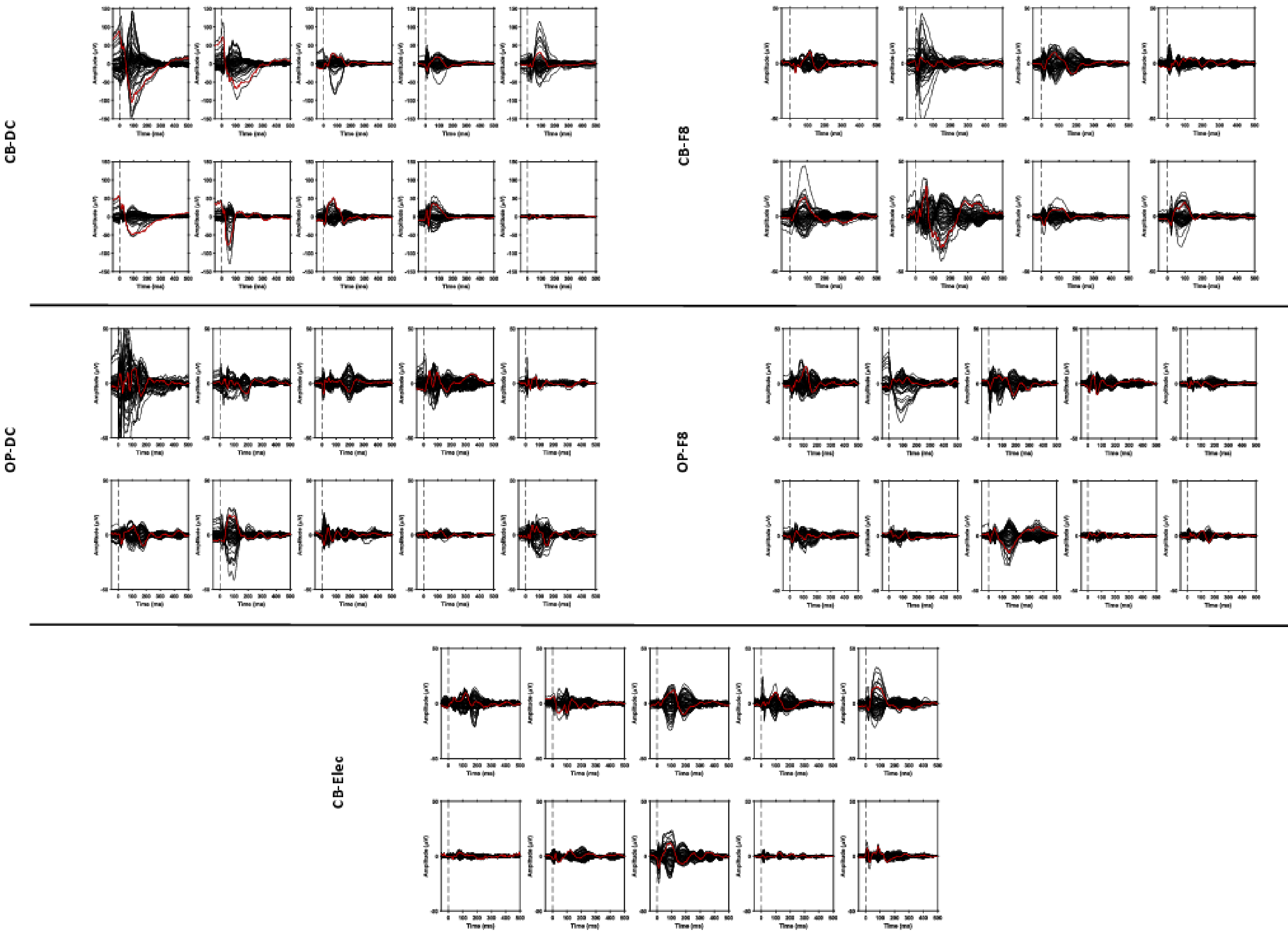


*Figure S3:* Timeseries for each participant following cleaning using the Original pipeline. Note the variability within conditions. Two participants were removed from the CB-F8 condition due to poor signal quality. Channel 'Oz' highlighted in red**.**

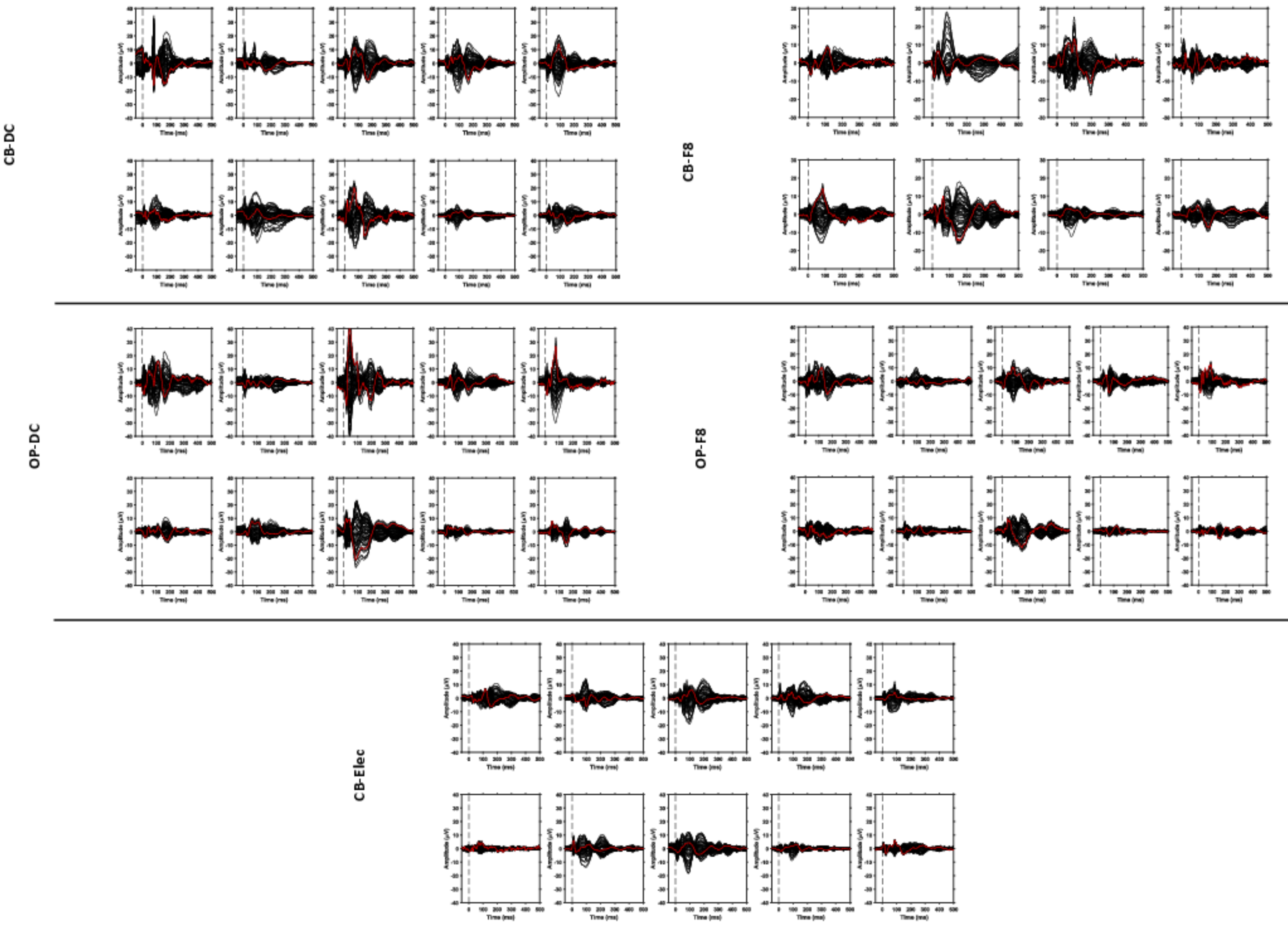


*Figure S4:* Timeseries for each participant following cleaning using the SOUND pipeline. Note the variability within conditions. Two participants were removed from the CB-F8 condition due to poor signal quality. Channel 'Oz' highlighted in red.

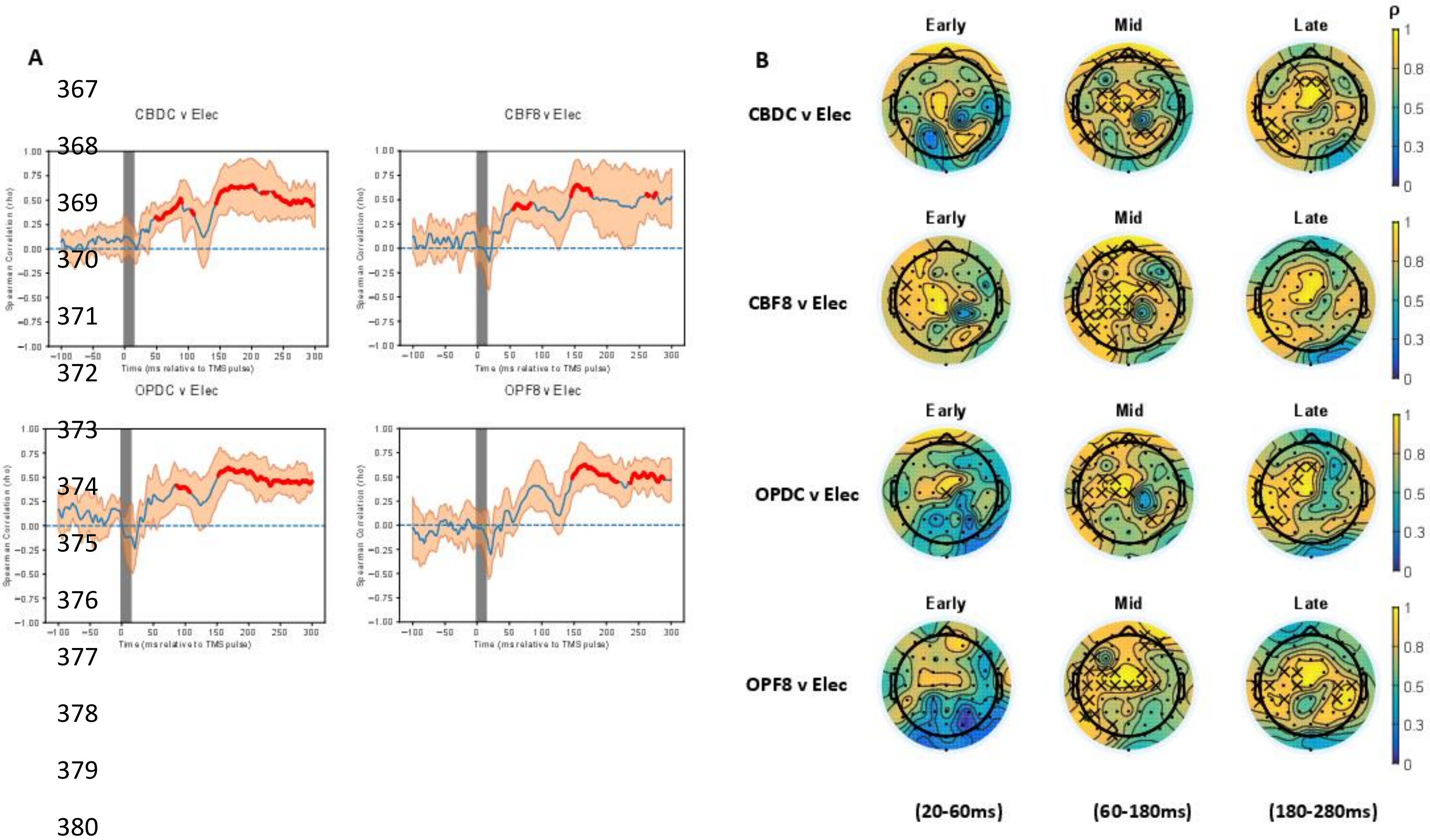


*Figure S5:* A. Spatial correlations following Laplacian filtering (SOUND pipeline). Red dots indicate significant positive correlations. No negative correlations were found. B. Temporal correlation analysis following Laplacian filtering for each active TMS condition against the electrical control following cleaning with the SOUND pipeline. Crosses indicate significant positive correlations. No negative correlations were found.

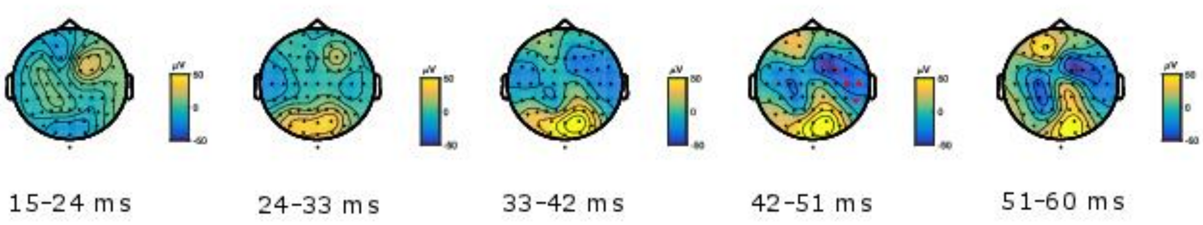


*Figure S6:* Cluster-based permutation tests comparing voltage distributions of peaks across OP-DC and CB-Electrical data cleaned with the SOUND pipeline and Laplacian filtering. Clusters defined as at least two adjacent electrodes exceeding p-value < 0.05 at each time-point. Critical alpha level for Monte-Carlo p-values set at p<0.025, 5000 iterations. Crosses indicate statistically significant different electrode clusters between conditions.

### *11. Supplementary Material – Additional Tables*

| | Trials Rejected | | Channels Rejected | |
|---|---|---|---|---|
| Condition | Orig. | SOUND | Orig. | SOUND |
| CB-DC | 0.04 ± 0.02 | 0.05 ± 0.06 | 0.002 ± 0.005 | 0.03 ± 0.03 |
| CB-Elec | 0.05 ± 0.03 | 0.04 ± 0.03 | 0.02 ± 0.03 | 0.03 ± 0.02 |
| CB-F8 | 0.06 ± 0.04 | 0.06 ± 0.04 | 0.01 ± 0.03 | 0.05 ± 0.03 |
| OP-DC | 0.06 ± 0.05 | 0.06 ± 0.04 | 0.007 ± 0.02 | 0.03 ± 0.03 |
| OP-F8 | 0.05 ± 0.02 | 0.05 ± 0.03 | 0.00 ± 0.00 | 0.05 ± 0.04 |

*Table S1:* Proportion (of total) of trials rejected and channels removed in the Original and SOUND pipelines for each condition. Shown as mean ± standard deviation.

| Scale | TMS Condition | Mean | SEM | SD |
|---|---|---|---|---|
| Discomfort | CB-DC | 4.45 | .45 | 1.42 |
| | CD-DS7A | 3.82 | .53 | 1.66 |
| | CB-F8 | 2.99 | .50 | 1.58 |
| | OP-DC | 3.6 | .43 | 1.35 |
| | OP-F8 | 2.6 | .64 | 2.01 |
| Pain | CB-DC | 2.32 | .31 | .97 |
| | CD-DS7A | 3.00 | .70 | 2.21 |
| | CB-F8 | 1.72 | .58 | 1.82 |
| | OP-DC | 1.62 | .44 | 1.41 |
| | OP-F8 | 1.42 | .42 | 1.33 |
| Twitch | CB-DC | 5.12 | .72 | 2.27 |
| | CD-DS7A | 3.62 | .77 | 2.43 |
| | CB-F8 | 3.47 | .66 | 2.09 |
| | OP-DC | 3.52 | .57 | 1.81 |
| | OP-F8 | 1.99 | .42 | 1.34 |
| Masking | CB-DC | 5.25 | .76 | 2.40 |
| | CD-DS7A | 5.77 | .60 | 1.88 |
| | CB-F8 | 6.25 | .72 | 2.28 |
| | OP-DC | 4.95 | .73 | 2.31 |
| | OP-F8 | 6.02 | .70 | 2.21 |

*Table S2:* Descriptive statistics for the VAS scales [0,10], for each TMS condition (site, coil type). CB - cerebellar site, OP - occipital/parietal site, DC - double-cone coil, DS7A – electrical (sham) stimulation, F8 – figure of 8 coil.

| Condition | [Min, Max] µV | Difference (Max-Min) µV |
|---|---|---|
| CB-DC | [-220880, 248830] | 469,710 |
| CB-Elec | [-215980, 131010] | 346,990 |
| CB-F8 | [-219100, 234800] | 453,900 |
| OP-DC | [-272870, 244950] | 517,820 |
| OP-F8 | [-166300, 234600] | 400,900 |

*Table S3*: Voltage deflections in the raw signals (min, max) for each condition in µV.

| Range Type | Condition | Mean | Median | SD | SEM |
|---|---|---|---|---|---|
| Voltage (µV) | CB-DC | 440900.00 | 440500.00 | 18799.82 | 5945.03 |
| | CB-Elec | 290000.00 | 290000.00 | 26495.28 | 8831.76 |
| | CB-F8 | 431625.00 | 437500.00 | 18631.29 | 6587.16 |
| | OP-DC | 450400.00 | 446000.00 | 22106.81 | 6990.79 |
| | OP-F8 | 332100.00 | 321500.00 | 53984.46 | 17071.39 |
| Recovery Time (ms) | CB-DC | 443.44 | 467.15 | 318.23 | 100.63 |
| | CB-Elec | 16.80 | 17.10 | 5.29 | 1.76 |
| | CB-F8 | 26.90 | 15.75 | 30.29 | 10.71 |
| | OP-DC | 62.43 | 40.25 | 57.47 | 18.17 |
| | OP-F8 | 109.80 | 18.70 | 279.87 | 88.50 |

*Table S4:* Descriptive statistics for peak voltage (µV) and recovery time (ms post-pulse) ranges (max – min) across all channels.

| | Voltage Range | | | Recovery Time Range | | |
|---|---|---|---|---|---|---|
| Contrast | *T* | *p* | *r* | *T* | *p* | *r* |
| CB-DC vs CB-Elec | .00 | .01 | -.63 | 1.00 | .01 | -.60 |
| CB-DC vs CB-F8 | 12.00 | .40 | -.21 | 2.00 | .03 | -.56 |
| CB-DC vs OP-DC | 36.5 | .36 | .21 | 2.00 | .01 | -.58 |
| CB-DC vs OP-F8 | .00 | .01 | -.63 | 10.00 | .07 | -.40 |
| CB-F8 vs OP-F8 | .00 | .01 | -.63 | 12.00 | .40 | -.21 |
| OP-DC vs OP-F8 | .00 | .01 | -.63 | 15.00 | .20 | -.28 |
| CB-F8 vs CB-Elec | .00 | .01 | -.63 | 14.00 | .58 | -.14 |
| OP-DC vs CB-Elec | 45.00 | .01 | .63 | 45.00 | .01 | .63 |
| OP-F8 vs CB-Elec | 35.00 | .14 | .35 | 31.00 | .31 | .24 |

*Table S5:* Wilcoxon signed rank tests assessing differences in peak voltage (µV) and recovery time (ms post-pulse) ranges (max – min) in different conditions. Taken across all channels.

| | Trials Rejected | | Channels Rejected | |
|---|---|---|---|---|
| Condition | Orig. | SOUND | Orig. | SOUND |
| CB-DC | 0.04 ± 0.02 | 0.05 ± 0.06 | 0.002 ± 0.005 | 0.03 ± 0.03 |
| CB-Elec | 0.05 ± 0.03 | 0.04 ± 0.03 | 0.02 ± 0.03 | 0.03 ± 0.02 |
| CB-F8 | 0.06 ± 0.04 | 0.06 ± 0.04 | 0.01 ± 0.03 | 0.05 ± 0.03 |
| OP-DC | 0.06 ± 0.05 | 0.06 ± 0.04 | 0.007 ± 0.02 | 0.03 ± 0.03 |
| OP-F8 | 0.05 ± 0.02 | 0.05 ± 0.03 | 0.00 ± 0.00 | 0.05 ± 0.04 |

*Table S6:* Proportion (of total) of trials rejected and channels removed in the Original and SOUND pipelines for each condition. Shown as mean ± standard deviation.

| Pipeline | Condition | [Min, Max] μV | Difference (Max-Min) μV |
|---|---|---|---|
| Original | CB-DC | [-154.10, 144.13] | 298.23 |
| | CB-Elec | [-24.87, 32.96] | 57.83 |
| | CB-F8 | [-57.06, 45.30] | 102.36 |
| | OP-DC | [-77.42, 60.51] | 137.93 |
| | OP-F8 | [-35.57, 26.40] | 61.97 |
| SOUND | CB-DC | [-24.41, 34.01] | 58.42 |
| | CB-Elec | [-19.21, 14.31] | 33.52 |
| | CB-F8 | [-17.78, 28.09] | 45.87 |
| | OP-DC | [-42.66, 52.62] | 95.28 |
| | OP-F8 | [-15.47, 15.76] | 31.23 |

*Table S7*: Voltage deflections (min, max) for each condition (μV) for both pipelines.

| | | Variance Removed (%) | | | | Components Removed (%) | | | |
|---|---|---|---|---|---|---|---|---|---|
| Stim. Cond. | Pipeline/ ICA Stage (1,2) | Mean | SEM | Median | SD | Mean | SEM | Median | SD |
| CB-DC | Original 1 | 46.85 | 6.31 | 38.88 | 19.96 | 0.12 | 0.01 | 0.12 | 0.03 |
| | 2 | 38.63 | 9.86 | 29.56 | 31.19 | 0.46 | 0.03 | 0.45 | 0.11 |
| | SOUND 1 | 45.2 | 4.74 | 45.06 | 14.97 | 0.12 | 0.01 | 0.11 | 0.03 |
| | 2 | 48.75 | 8.67 | 41.10 | 27.41 | 0.27 | 0.03 | 0.29 | 0.10 |
| Elec | Original 1 | 37.63 | 8.33 | 30.93 | 26.34 | 0.04 | 0.01 | 0.04 | 0.02 |
| | 2 | 37.35 | 10.04 | 29.93 | 31.74 | 0.43 | 0.03 | 0.43 | 0.08 |
| | SOUND 1 | 28.61 | 4.59 | 30.33 | 14.52 | 0.05 | 0.00 | 0.05 | 0.01 |
| | 2 | 11.85 | 4.30 | 3.18 | 13.58 | 0.19 | 0.02 | 0.17 | 0.05 |
| CB-F8 | Original 1 | 41.94 | 6.41 | 43.23 | 20.27 | 0.10 | 0.01 | 0.10 | 0.03 |
| | 2 | 29.77 | 8.15 | 23.54 | 25.77 | 0.47 | 0.02 | 0.48 | 0.07 |
| | SOUND 1 | 33.64 | 4.14 | 28.78 | 13.1 | 0.09 | 0.01 | 0.09 | 0.03 |
| | 2 | 23.69 | 8.87 | 19.37 | 28.04 | 0.24 | 0.03 | 0.22 | 0.10 |
| OP-DC | Original 1 | 57.07 | 6.16 | 58.88 | 19.48 | 0.12 | 0.01 | 0.13 | 0.04 |
| | 2 | 58.02 | 9.88 | 67.09 | 31.24 | 0.48 | 0.04 | 0.46 | 0.11 |
| | SOUND 1 | 37.38 | 4.25 | 38.26 | 13.43 | 0.10 | 0.01 | 0.09 | 0.04 |
| | 2 | 29.67 | 8.57 | 27.15 | 27.11 | 0.23 | 0.03 | 0.24 | 0.09 |
| OP-F8 | Original 1 | 66.66 | 7.52 | 70.98 | 23.79 | 0.08 | 0.01 | 0.07 | 0.03 |
| | 2 | 44.47 | 6.41 | 40.95 | 20.25 | 0.51 | 0.02 | 0.48 | 0.08 |
| | SOUND 1 | 38.70 | 6.06 | 34.04 | 19.15 | 0.08 | 0.01 | 0.09 | 0.02 |
| | 2 | 29.42 | 6.62 | 20.26 | 20.93 | 0.29 | 0.03 | 0.29 | 0.09 |

*Table S8:* Descriptive statistics for two stages of ICA output – % variance removed and % components removed.

| | Variance Removed (%) | | | Components Removed (%) | | |
|---|---|---|---|---|---|---|
| Cond. Contrast | T | p | r | T | p | r |
| CB-DC | 28.00, 36.00 | .96, .39 | .01, .19 | 28.00, 0.00 | .96, .01 | .01, -.63 |
| CB-Elec | 13.00, 9.00 | .14, .06 | -.33, -.42 | 47.00, 0.00 | .05, .01 | .44, -.63 |
| CB-F8 | 18.00, 23.00 | .33, .65 | -.22, -.10 | 16.00, 0.00 | .44, .01 | -.17, -.63 |
| OP-DC | 3.00, 7.00 | .01, .04 | -.56, -.47 | 12.00, 0.00 | .11, .01 | -.35, -.63 |
| OP-F8 | 1.00, 14.00 | .01, .17 | -.60, -.31 | 40.00, 0.00 | .20, .01 | .29, -.63 |

*Table S9:* Wilcoxon Signed Rank comparisons between Original and SOUND pipelines for ICA output (ICA step 1, ICA step 2).
